# Alzheimer’s disease-associated β-Amyloid does not protect against Herpes Simplex Virus 1 brain infection

**DOI:** 10.1101/2021.01.21.427570

**Authors:** Olga Bocharova, Kara Molesworth, Narayan P. Pandit, Ilia V. Baskakov

## Abstract

Alzheimer’s disease (AD) is devastating fatal neurodegenerative disease. An alternative to the amyloid cascade hypothesis is the hypothesis that a viral infection is key to the etiology of late-onset AD, with amyloid Aβ peptides playing a protective role. Contrary to previous work, in the current study the 5XFAD genotype failed to protect mice against infection with two strains of herpes simplex virus 1 (HSV-1), 17syn+ and McKrae. Moreover, the region- or cell-specific tropisms of HSV-1 were not affected by the 5XFAD genotype, arguing that host-pathogen interactions were not altered. In aged 5XFAD mice with abundant Aβ plaques, only small, statistically non-significant protection against acute HSV-1 infection was observed, yet no colocalization between HSV-1 and Aβ plaques was found. While the current study questions the antiviral role of APP or Aβ, it neither supports nor refutes the viral etiology hypothesis of late-onset AD.

## Introduction

Alzheimer’s disease (AD) is devastating fatal neurodegenerative disease, which is estimated to affect 5.8 million Americans in 2020. The vast majority of AD cases are late-onset, which is believed to be sporadic in origin. Over the years, a number of gene variants that increase the susceptibility to late-onset AD have been identified [1], yet our understanding of the etiology of the disease still remains poor. According to the amyloid cascade hypothesis, the pathological cascade of events leading to AD is triggered by aggregation and accumulation of Aβ peptides 1-40 and 1-42, the proteolytic fragments of Amyloid Precursor Protein (APP) [2, 3].

An alternative to the amyloid cascade hypothesis is the hypothesis that a viral or microbial infection of the CNS is an essential component of the etiology of late-onset AD [4–6]. According to this hypothesis, which is known as the antimicrobial or antiviral protection hypothesis, Aβ peptides possess antimicrobial and antiviral effects and are produced by the CNS as a defense mechanism [7–10]. Viral and microbial pathogens trigger a pathological cascade, presumably by seeding of Aβ fibrils or plaques, which can entrap and neutralize CNS pathogens [7, 9, 11]. While individuals who experienced a single viral challenge or have a latent infection might not be at a high risk for late-onset AD, recurrent reactivation of a latent infection, or an accumulative effect of low-grade chronic infection, and multiple viral challenges over the life-time places an individual at high-risk for late-onset AD. Such risk is realized in individuals expressing the high-risk AD variants of *APOE, TREM2, CR1, CD33* and other genes, many of which are associated with innate immunity. A number of pathogens including *Chlamydia pneumonia* [12–15], *Borrelia spirochetes* [16, 17], Herpes zoster [18], human Herpesviruses 6 and 7 (HHV6 and HHV7) [19] and Herpes Simplex Virus 1 (HSV-1) [4, 20, 21] have been linked to the late-onset AD over the years. Among the broad range of pathogens, HSV-1 has emerged as one of the leading pathogens linked to late-onset AD in a number of independent studies (reviewed in [4, 22, 23]).

A recent study by Dudley and coauthors examined hundreds of brains across multiple datasets and reported a greater abundance of HHV6 or HHV7 RNA and DNA in brains of late-onset AD individuals relative to controls [24]. This study suggested that herpes viruses drive the production of Aβ peptides [24]. Simultaneously, the work by Eimer and coauthors showed that Aβ peptides protect neurons in 3D cultures and prolong the survival of young 5XFAD mice infected with HSV-1 [7]. A recent study reanalyzed the data from Dudley’s work and concluded that the statistical analysis in Dudley’s study was too weak to prove a link between viral load and AD [25]. Moreover, the most recent work by Jacobson and coauthors showed no differences between postmortem AD and control human brains with respect to viral RNA/DNA load [26]. While Jacobson’s study questioned previous results, a viral role in the etiology of AD was not ruled out by the new results.

Upon isolation of multiple strains of HSV-1 (strain is defined as plaque-purified clinical isolate [27]), a genomic diversity of HSV-1 has been demonstrated [28]. HSV-1 strain-specific features were shown to dictate important aspects of host-pathogen interaction including the median lethal dose (LD_50_) value, reactivation from latency and, possibly, even cell tropism [27, 29–31]. In order to examine whether the protective role of 5XFAD genotype is dictated by the strain identity of HSV-1, in the current study we employed the approach introduced by Eimer and coauthors [7]. This approach involves testing 5XFAD mice that overexpress mutant human APP with the Swedish (K670N, M671L), Florida (I716V), and London (V717I) familial AD mutations along with human presenilin 1 (PS1) harboring two mutations, M146L and L286V [32] to resist acute HSV-1 infection. Surprisingly, contrary to previous results [7], in the current work the 5XFAD genotype failed to protect the mice upon challenges with two HSV-1 strains, 17syn+ and McKrae. Moreover, the region-specific or cell-specific tropisms of HSV-1 strains were not affected by 5XFAD genotype, when compared to wild type (WT) littermate controls, suggesting that the host-pathogen interactions were not affected by APP overexpression. Aged 5XFAD animals, in which Aβ plaques were abundant, showed a delay and slightly better survival rate relative to WT mice, yet the differences between the two groups were not statistically significant. Moreover, in aged 5XFAD mice, Aβ plaques were free of HSV-1 viral particles, suggesting that preformed plaques do not entrap the virus. In summary, the current results question an antiviral role for APP or Aβ. Nevertheless, the current work neither supports nor refutes the hypothesis of the viral etiology of late-onset AD.

## Materials and Methods

### Key Resource Table

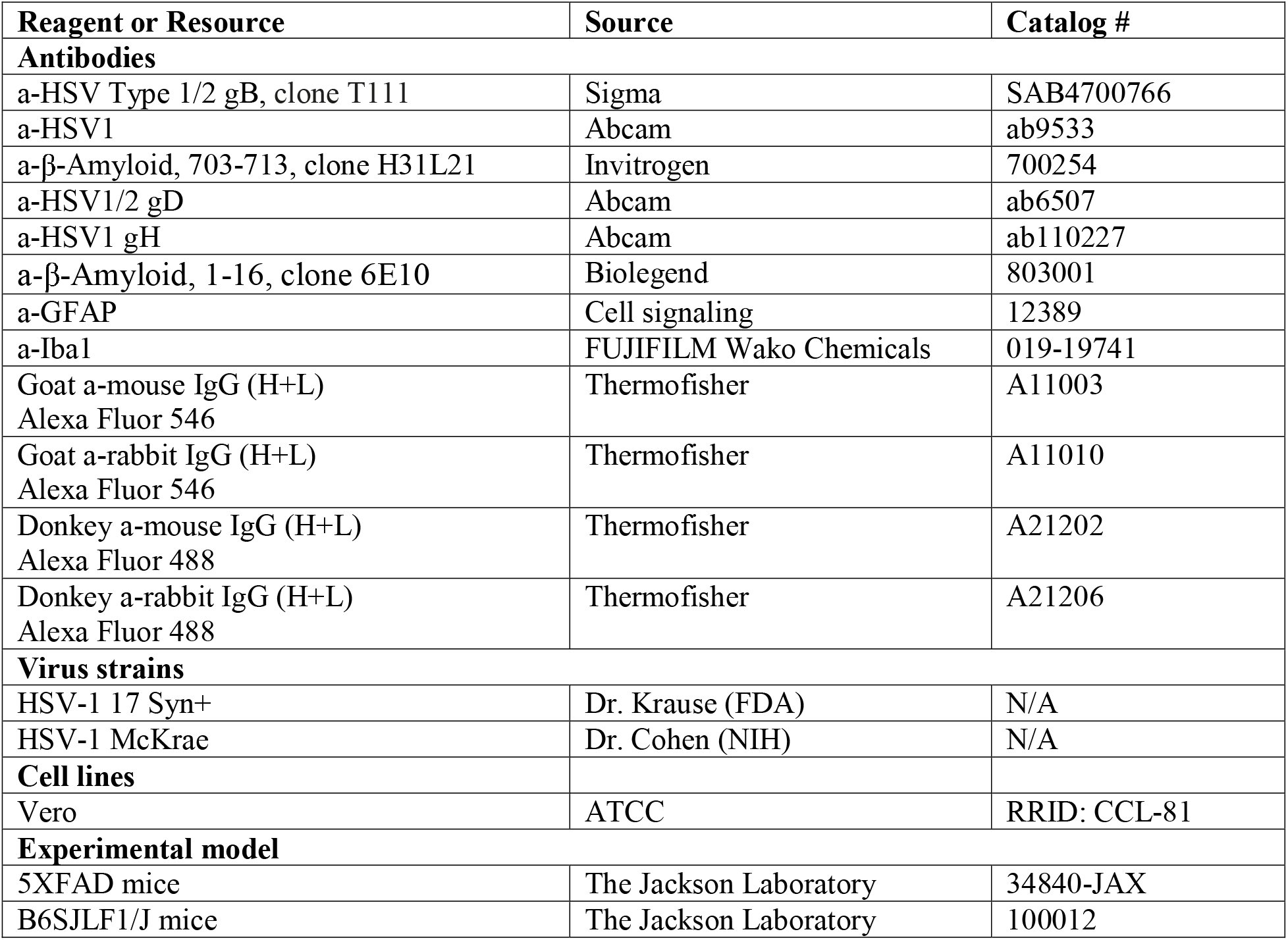

### Experimental models

Male and female 5-6 week old 5XFAD mice [B6SJL-Tg(APPSwFlLon, PSEN1*M146L*L286V) 6799Vas/Mmjax] and WT littermates were used for inoculation experiments. 5XFAD and WT littermates were group-housed together in random ratios, three to five mice per cage, and kept on a 12 hour light/dark cycle. The ratios of male and female for both, 5XFAD and WT littermates, in each experiment were random. The study was carried out in strict accordance with the recommendations in the Guide for the Care and Use of Laboratory Animals of the National Institutes of Health. The animal protocol was approved by the Institutional Animal Care and Use Committee of the University of Maryland, Baltimore (Assurance Number: A32000-01; Permit Number: 0419007).

### Production and titration of HSV-1 stocks

HSV-1 17 Syn+ and McKrae strains were propagated in Vero cell culture with initial multiplicity of infection (MOI) of 0.01 PFU/cell in serum-free medium. After 2 hours of incubation at 37°C the viral inoculum was aspirated, and the cell culture was supplemented with fresh DMEM medium with 5% newborn calf serum. Cells were incubated for 2 to 3 days, until 100% of the cells displayed cytopathic effects. After one freezing and thawing cycle cells were sonicated 5 times for 30 sec using a Misonix S-4000 sonicator at 600 W output, letting the cell suspension cool on ice for 1 min between sonications. After centrifugation of the Vero cell lysate at 12000 x g for 10 min at 4°C, virus-containing supernatant was mixed with 10% sterile BSA solution to a final concentration of 2% BSA, and supplemented with 10x PBS to a final concentration of 1x PBS (with 137 mM of NaCl). Viral stock was split into smaller aliquots in screw-capped cryovials and stored at −80°C. The HSV-1 17 Syn+ strain was a generous gift of Dr. Krause (FDA) and HSV-1 McKrae strain was a generous gift of Dr. Cohen (NIH).

Viral titer was measured by standard plaque assay on Vero cell culture under 1.8% agar layer. 10-fold serial dilutions in 1 ml serum-free medium were added to the corresponding well of 6-well plate in duplicates and incubated for 2 hours with swirling every 30 min. Plates underwent plaque count after 2-3 days of incubation. Vero monolayer was rinsed twice with warm PBS. One ml methanol was added to each well and left flat for 5 min at room temperature to fix cells. After aspiration of the methanol 0.5 ml of 0.5% Crystal Violet in 25% methanol was added to visualize plaques. To determine titer of the stock, number of plaques in duplicate was averaged for a given dilution.

### Bioassay

5XFAD APP/PS1 transgenic mice (Oakley et al., 2006) that overexpress Aβ42 from inheritable FAD mutant forms of human APP (the Swedish mutation: K670N, M671L; the Florida mutation: I716V; the London mutation: V717I) and PS1 (M146L; L286V) transgenes under the transcriptional control of the neuron-specific mouse Thy1 promoter (Tg6799 line). 5XFAD lines (with B6SJLF1/J genetic background) were purchased from Jackson Laboratory and maintained by crossing heterozygous 5XFAD transgenic males with B6SJLF1/J female breeders. All pups were genotyped using Transnetyx genotypic services (Cordova, TN, USA). All 5XFAD transgenic mice were heterozygotes with respect to the transgene. The ratios of males to females for both 5XFAD and WT littermates in each experiment were random. 5XFAD and WT littermates were group-housed together in random ratios, three to five mice per cage for the entire experiment. Control animals were caged separately from the animals inoculated with HSV-1. 17Syn+ and McKrae inoculation stocks were prepared in PBS containing 137 mM of NaCl and 2% sterile BSA and supplemented with antibiotics-antimycotics concentrate (Invitrogen). Immediately before inoculation, a stock was diluted with PBS-NaCl-BSA sterile solution to the required titer and inoculated into 5-6 week-old mice. Each 5XFAD or WT littermate control mouse received a single 20 μl of inoculum intracerebrally (IC) under 2% isoflurane anesthesia, in the center of left hemisphere 2 mm from the sagittal suture. The injection depth of 2 mm was achieved using a needle length stopper made from a needle cap. After inoculation, animals were observed and weighed daily for the following signs of acute Herpes Simplex virus encephalitis: eye and face inflammation, walking difficulty, tremors, hunched posture, and weight loss. The animals were euthanized when they demonstrate the severe aforementioned clinical signs and/or 20% loss of their weight, as determined relative to weights measured prior to virus inoculation.

### Histopathology and Immunofluorescence

After euthanasia by CO_2_ asphyxiation, brains were immediately extracted and kept ice-cold during dissection. Brains were sliced using a rodent brain slicer matrix (Zivic Instruments, Pittsburg, PA). Three◻mm central coronal sections of each brain were formalin-fixed and embedded in paraffin. Four◻μm sections were produced using a Leica RM2235 microtome (Leica Biosystems, Buffalo Grove, IL) were mounted on slides and processed for immunohistochemistry.

Rehydrated slides were subjected to epitope exposure for 20◻min of hydrated autoclaving at 121°C in trisodium citrate buffer, pH◻6.0, with 0.05% Tween 20. All antibodies used in this study are listed in the Key Resources Table. Immunofluorescence detection was performed by using AlexaFluor-488 and AlexaFluor-546 labeled secondary antibodies. An autofluorescence eliminator (Sigma-Aldrich, St. Louis, MO) was used according to the original protocol to reduce background fluorescence. Fluorescent images were collected using an inverted Nikon Eclipse TE2000-U microscope (Nikon Instruments Inc, Melville, NY) equipped with an illumination system X-cite 120 (EXFO Photonics Solutions Inc., Exton, PA, United States) and a cooled 12-bit CoolSnap HQ CCD camera (Photometrics, Tucson, AZ, United States). Images were processed using WCIF ImageJ software (National Institute of Health, Bethesda, MD, United States).

### Analysis of viral genome number

Frontal and rear brain parts remaining after a central 3 mm coronal cut were separated into right and left halves and kept frozen at −80°C for subsequent DNA isolation. For analysis of viral copy number, the left frontal portions of brains were used. DNA isolation was performed according to the manufacturer’s instructions using QIAamp DNAMiniKit (QIAGEN, Germany). Copy numbers of HSV-1 were measured using Virusys Corporation’s HSV-1 qPCR kit with a FAM/BHQ-labeled probe specific for glycoprotein D (gD) of HSV-1 (Virusys, Taneytown, MD, USA, cat. # H1K240). To build a calibration curve for determining an absolute copy number, serial 1/10 dilutions of the Virusys kit standard were made that produced linear dependence within a range of 10^2^ to 10^8^ copies per reaction. PCR product was detected by CFX96 Real-Time PCR Detection System (Bio-Rad, Hercules, CA, USA).

### Quantification of Aβ plaques and HSV-1 replication centers

Images were taken under 10x magnification and 500 ms exposure time. For quantification of plaques and HSV-1 replication centers, the images taken in green (anti-Aβ) and red (anti-HSV-1) channels were converted to 8-bit greyscale and then subjected to ImageJ signal area measurement after automated thresholding. On images taken in the red channel, non-specific spots arising from blood vessels were deleted prior to analysis procedure.

### Quantification and statistical analysis

Statistical analysis of survival curves was performed using GraphPad Prism software, versions 8.4.2 for Windows (GraphPad Software, San Diego, California USA). Survival curves were compared using Log-rank (Mantel-Cox) test. Differences with the p values > 0.05 were considered to lack statistical significance.

Statistical analysis of integrated density of Aβ plaques and HSV-1 shown in Figure 6C was performed using GraphPad Prism software, versions 8.4.2 for Windows (GraphPad Software, San Diego, California USA). The differences in integrated density of Aβ and HSV-1 signals in WT and 5XFAD brains were analyzed by ordinary 2-way ANOVA with Šidák’s multiple comparisons test.

Statistical analyses of plaques and virus colocalization were performed using Colocalization Threshold plugin of ImageJ software (National Institute of Health, Bethesda, MD, United States). After determination of threshold by the Costes method [33], the thresholded Mander’s Split Colocalization coefficients were calculated for each channel (red and green). The box-and-whisker plot of tM1 and tM2 values in Figure S5 was built using Excel. The midline denotes the median, the x represents the mean and the ends of the box-and-whisker plot denote the 25th and 75th percentiles. The whiskers extend from the ends of the box to the minimum value and maximum value. The interquartile range (IQR) was defined as the distance between the 1^st^ quartile and the 3^rd^ quartile. A data point was considered an outlier if it exceeded a distance of 1.5 times the IQR below the 1^st^ quartile or 1.5 times the IQR above the 3^rd^ quartile.

## Results

### Lack of Protective Effect of the 5XFAD Genotype in Young Mice

For preparing HSV-1 inoculation stocks, two commonly used HSV-1 strains, 17syn+ and McKrae, were propagated in Vero cells, and the resulting cell lysates were titrated using the same cell line. The McKrae strain was shown to be more neurovirulent and has a lower LD50 value relative to 17syn+ [27, 31, 34]. The HSV-1 dose is expressed in Plaque-Forming Units (PFUs) administered per animal. To test whether the 5XFAD genotype has protective effects against HSV-1 encephalitis, we used male and female 5XFAD and littermate B6SJL (WT) control mice of the same age as in previous studies [7], i.e. five- to six-week-old. Two strains of HSV-1, 17syn+ and McKrae, were administered intracranially (IC) to examine the strain-specificity of the effects. Three doses of each strain were tested, where the highest and the lowest doses were above and below of the LD_50_ values, respectively. As expected, the highest doses resulted in the highest mortality rates (Figures 1A and 1B). Reducing the viral dose increased the survival rates and prolonged the incubation times of non-survivors in both 5XFAD and WT cohorts challenged with both HSV-1 strains (Figures 1A and 1B). Out of six experimental conditions tested, 5XFAD mice showed higher survival rates relative to the WT littermates only in one condition: animals challenged with 10^4^ PFUs of McKrae (Figure 1B, upper panel), however, this difference between 5XFAD and WT cohorts lacked statistical significance. Moreover, lowering the inoculation dose two-fold, from 10^4^ PFUs to 5×10^3^ PFUs (Figure 1B, middle panel), reversed the survival yield between 5XFAD and WT littermates suggesting that minor variations in survival yield might be due to stochastic variations between experiments.

**Figure 1.**
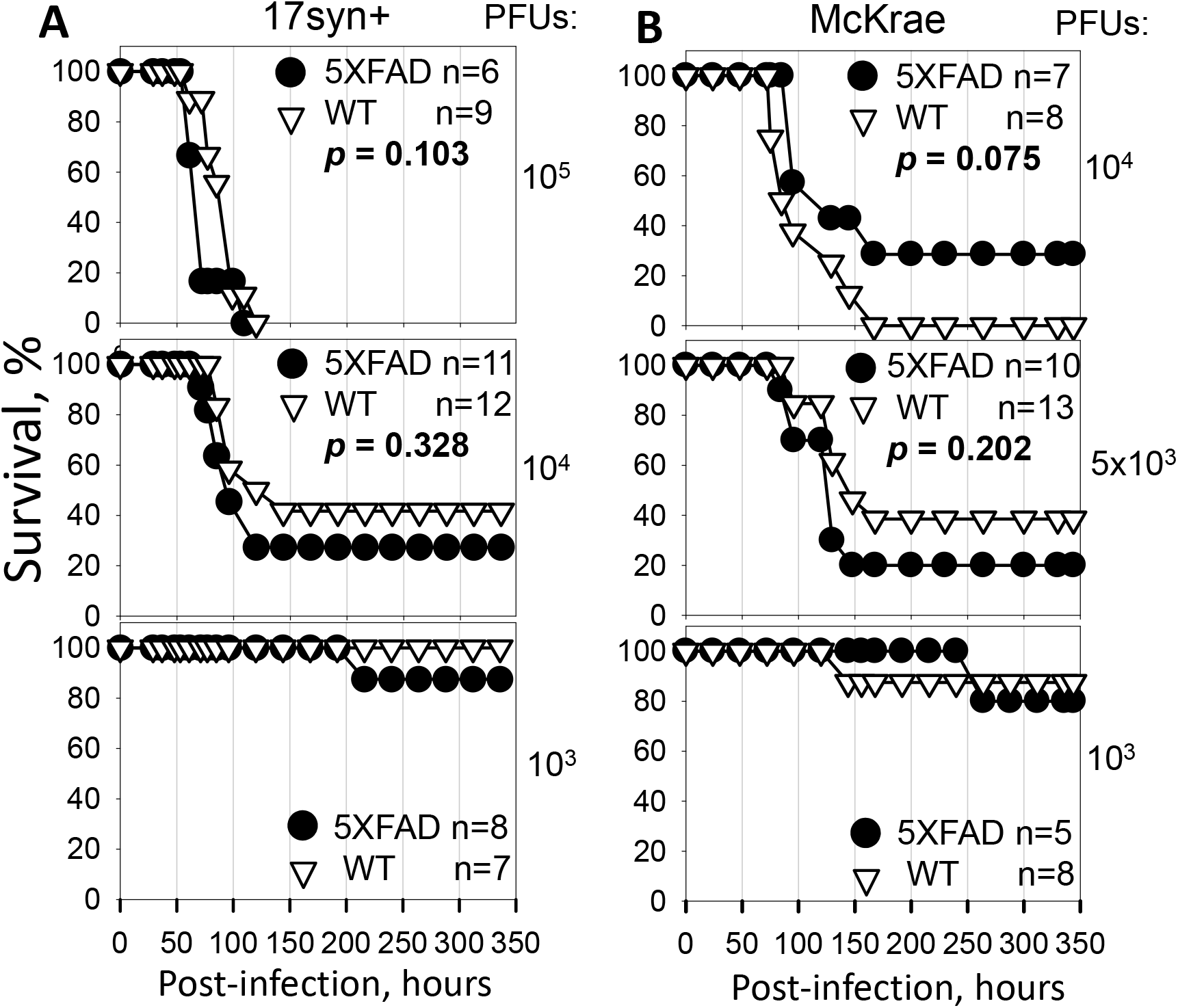
Dose-Response of Young 5XFAD Mouse Model to HSV-1 challenge. Survival curves for 5-6 week old 5XFAD and wild-type littermate (WT) mice challenged via IC injections with 10^5^, 10^4^ or 10^3^ PFUs of 17syn+ strain per mouse (**A**); or 10^4^, 5×10^3^, or 10^3^ PFUs of McKrae strain per mouse (**B**). 5XFAD and WT littermate mice were caged together in random ratios. Individual plots show independent experiments with number (*n*) of animals of each genotype indicated. Statistical significance (*p*) was calculated using log-rank (Mantel-Cox).

Because a previous study that documented protective effects of the 5XFAD genotype employed only female mice [7], the survival curves were re-analyzed for females only. Survival of females showed the same pattern as survival of males and females combined (Figures S1A and S1B) and, again, showed no statistically significant differences between 5XFAD and WT cohorts. In summary, these experiments employing two HSV-1 strains, 17syn+ and McKrae, revealed a lack of protective effect of the 5XFAD genotype against HSV-1 encephalitis.

### Region-Specific Tropism of HSV-1 Is Not Altered in 5XFAD Mice

To test whether high expression levels of the disease-associated APP variant in 5XFAD mice alters HSV-1 tropism, brains of 5XFAD mice and WT littermates were analyzed using immunostaining for HSV-1. Infected cells were detected using a-HSV1 antibody against that stains HSV-1 replication centers [35]. Depending on replication stage, a-HSV1 staining displays granular or diffuse staining patterns within nuclei of infected cells (Figure 2) [35]. Hippocampus, hypothalamus, cortex and amygdala were consistently found to be among the most severely infected brain regions in both, 5XFAD and WT controls (Figures 2 and S2). No a-HSV1 staining was found in age-matched non-infected 5XFAD mice (Figure S2). While variations in regional distribution of HSV-1replication sites were found within animal groups, the most affected brain regions remained the same in animals of both genotypes.

**Figure 2.**
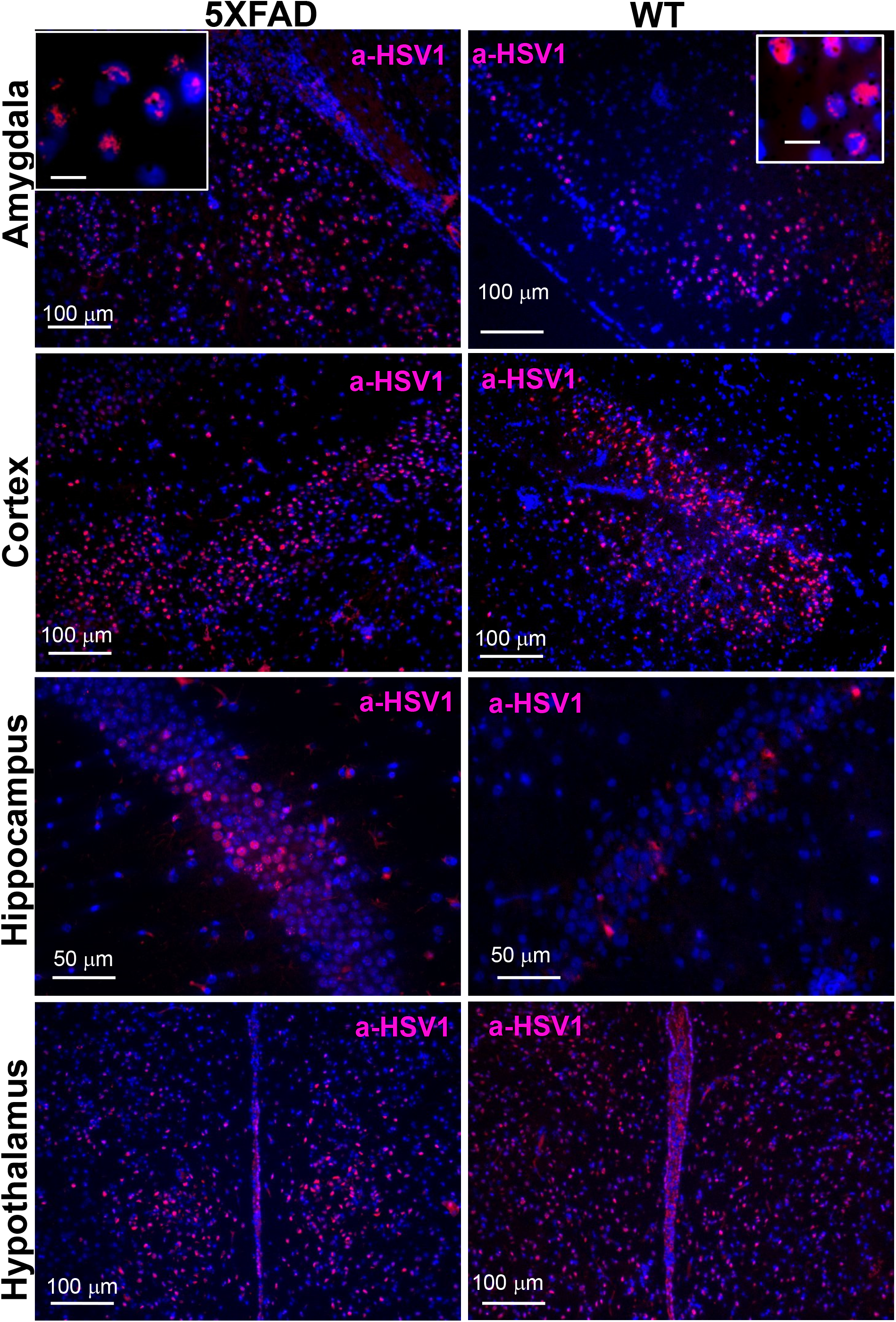
Region-Specific Tropism of HSV-1 Is Not Altered in Young 5XFAD Mice. Immunostaining for HSV-1 replication centers (a-HSV1 antibody, red) and cell nuclei (DAPI, blue) in 6-week old 5XFAD (*n*=6) and WT littermates (*n*=7) that did not survive IC challenges with 10^5^ PFUs of 17syn+.

### Cell-Specific Tropism of HSV-1 Is Not Altered in 5XFAD Mice

*In vitro,* HSV-1 infects different cell types including astrocytes, microglia and oligodendrocytes, whereas *in vivo* the virus predominantly invades neurons [36–40]. To test whether high expression of the human APP variant in neurons protects neuronal cells from invasion and alters cell tropism, brain slices were co-immunostained for HSV-1 replication sites and for glial fibrillary acidic protein (GFAP), a marker for astrocytes, or for Iba1, a microglia-specific marker. In both genotypes, 5XFAD and WT littermate controls, the vast majority of HSV-1-infected cells were neurons. Extremely rare HSV-1-infected astrocytes, and no HSV-1-positive microglia were found (Figures 3A and 3B). In both the 5XFAD and WT cohorts, reactive microglia were often found in close vicinity to HSV-1-infected neurons or were engulfing infected cells (Figures 3B). To summarize, no differences with respect to the infected cell types or infected brain areas were observed between 5XFAD and WT littermates.

**Figure 3.**
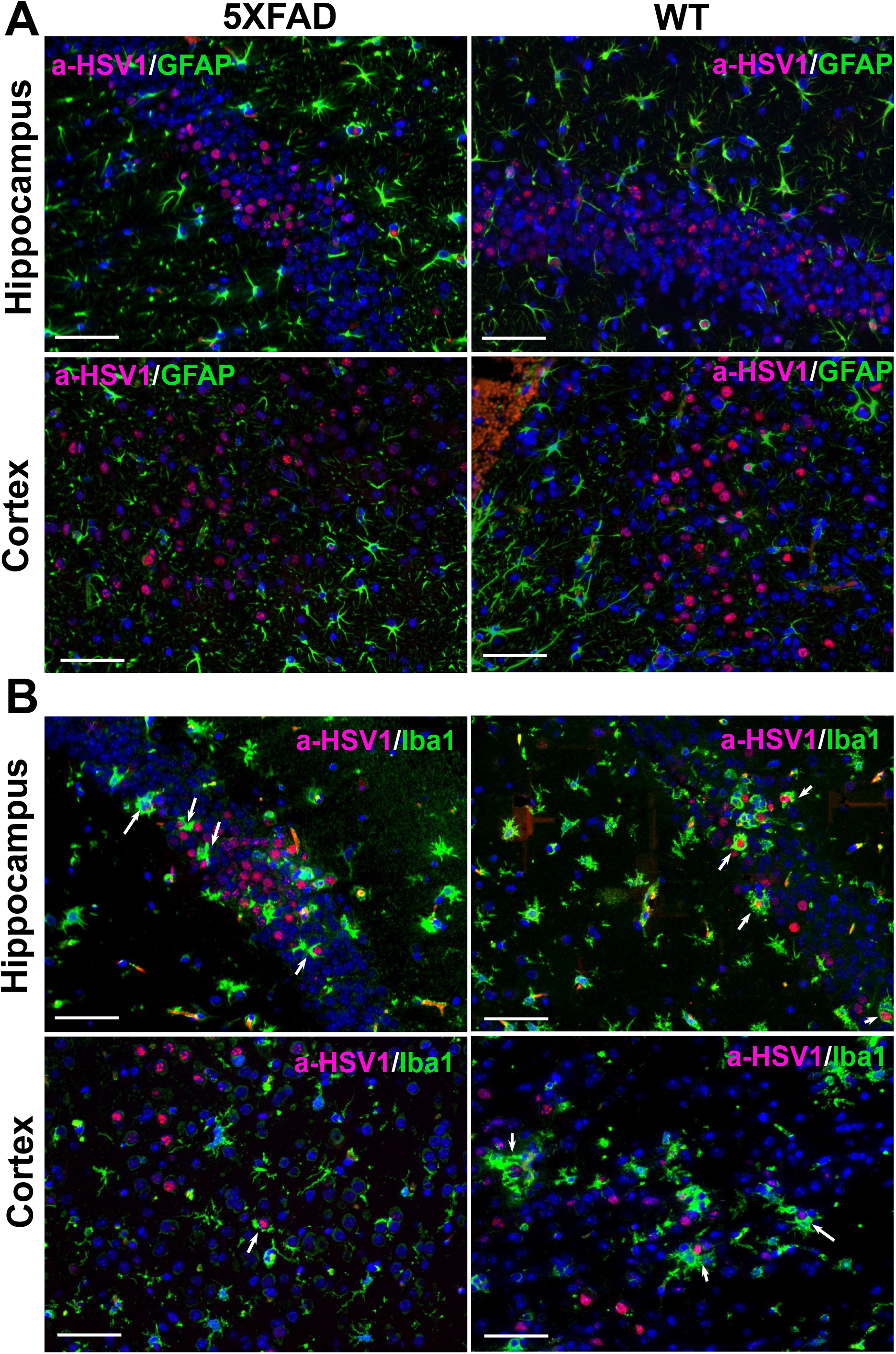
Cell-Specific Tropism of HSV-1 Is Not Altered in Young 5XFAD Mice. **(A)** Co-immunostaining for HSV-1 replication centers (a-HSV1 antibody, red), astrocytes (anti-GFAP antibody, green) and cell nuclei (DAPI, blue) in 6-week old 5XFAD (*n*=5) and WT littermates (*n*=5) that did not survive IC challenges with 10^4^ PFUs of 17syn+. **(B)** Co-immunostaining for HSV-1 replication centers (a-HSV1 antibody, red), microglia (anti-Iba1 antibody, green) and nuclei (DAPI, blue) in 5XFAD (*n*=5) and WT littermates (*n*=5) that did not survive IC challenges with 10^4^ PFUs of 17syn+. Arrows point at microglia engulfing HSV-1-infected cells. Scale bars = 50 μm.

### HSV-1 Infects Both APP-Positive and Negative Neurons

Next, we examined whether cells expressing high levels of APP are protected from HSV-1 invasion. In 5XFAD mice, APP is expressed under the Thy1 promoter and only a subpopulation of cells with active Thy1 promoter express high levels of APP. Under low magnification, the sites of active HSV-1 replication seemed to be excluded from the areas expressing high levels of APP (Figure 4A). It is not clear whether this effect was due to actual antiviral effects of APP or intrinsic tropism of HSV-1 to a subpopulation of neurons with low Thy1 activity. High magnification imaging revealed HSV-1 replication sites in both APP-positive and negative neurons (Figure 4B). This result suggests the possibility that the infection rate might be reduced by APP overexpression, yet APP does not fully protect cells from HSV-1 invasion and replication.

**Figure 4.**
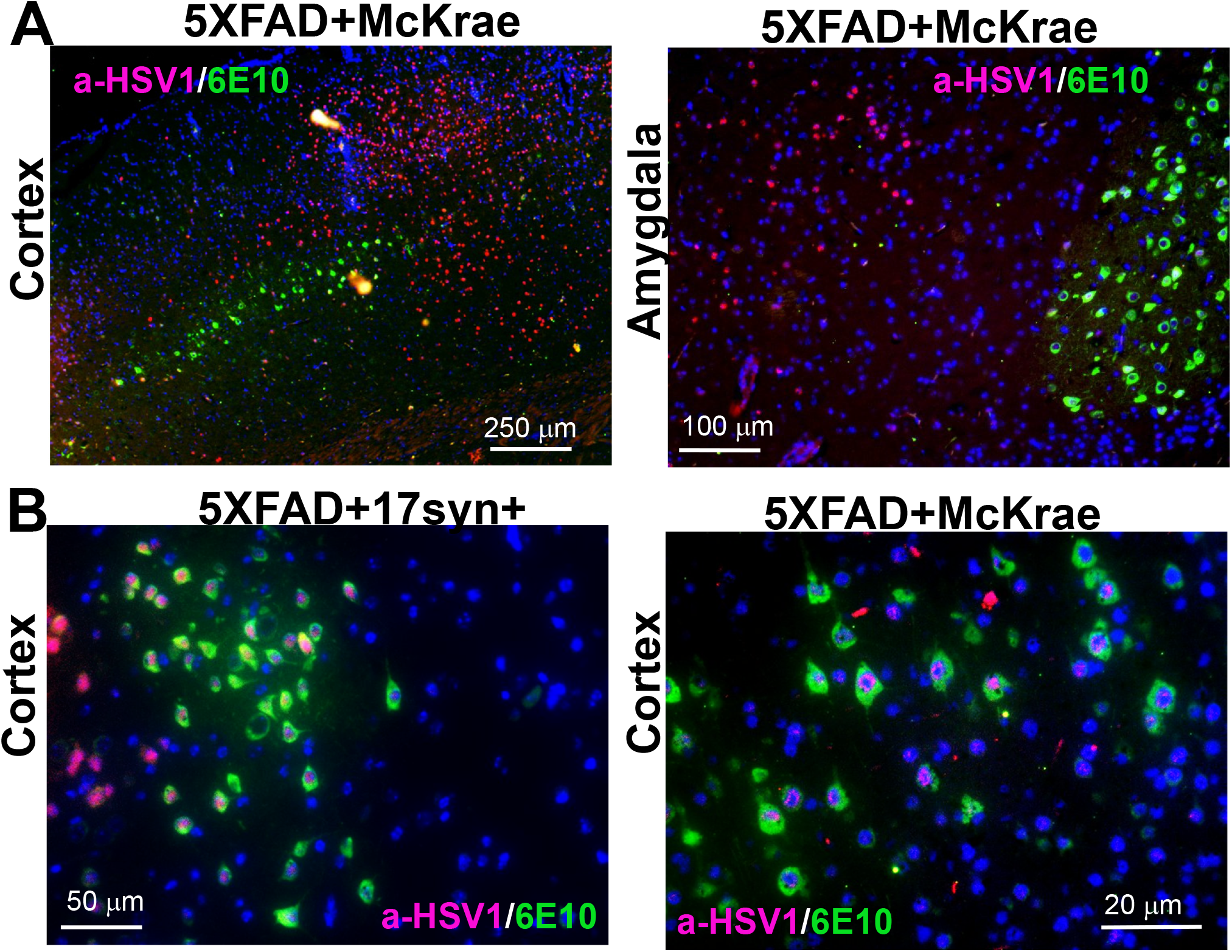
HSV-1 Infects Both APP-Positive and Negative Neurons. **(A)** Co-immunostaining for HSV-1 replication centers (a-HSV1 antibody, red), APP (6E10 antibody, green) and nuclei (DAPI, blue) in 6-week old 5XFAD (*n*=6) challenged IC with 5×10^3^ PFUs of McKrae. **(B)** Co-immunostaining for HSV-1 replication centers (a-HSV1 antibody, red), APP (6E10 antibody, green) and nuclei (DAPI, blue) in 5XFAD challenged IC with 10^5^ PFUs of 17syn+ (left panel, *n*= 2) or 10^4^ PFUs of McKrae (right panel, *n*=5).

### HSV-1 Does Not Induce Formation of Aβ Plaques in Young 5XFAD Mice

For testing whether HSV-1 triggers formation of Aβ plaques, 5XFAD mice that survived IC challenges with 17syn+ or McKrae were examined by co-immunostaining for Aβ and HSV-1 replication centers. No Aβ plaques or HSV-1 replication centers were detected in 5XFAD survivals 2 or 7.5 weeks post-inoculation (Figure S3A and S3B). Non-inoculated, aged 5XFAD mice, used as positive controls of Aβ staining, showed Aβ plaques (Figure S3C). Notably, in 5XFAD surviving mice, no Aβ plaques were found, including within those brain areas (motor cortex and hippocampus), where HSV-1 normally replicates under acute infection (Figure 2) and Aβ plaques form during aging (Figure S3C). These results suggest, that HSV-1 was cleared in 5XFAD survivors without triggering Aβ deposition.

### Lack of Protective Effect of Aβ in Aged 5XFAD Mice

Previous studies suggested that the antiviral effect of Aβ involves binding and entrapping of HSV-1 by Aβ peptides [7]. We reasoned that Aβ plaques could be more effective than non-aggregated Aβ in trapping viruses. In 5XFAD mice, the first Aβ plaques are found as early as four months of age. To test whether existing Aβ plaques are a prerequisite for the antiviral effect, 7 to 10-month old 5XFAD and WT littermates were challenged IC with the McKrae strain. Combined male and female 5XFAD mice showed a slightly better overall survival rate and, particularly, a better survival rate within the first 140 hours post-infection relative to WT littermates (Figure 5A). However, the difference between the 5XFAD and WT cohorts lacked statistical significance. Survival curves for females only showed the same trend as survival of combined males and females, yet also lacked statistically significant difference between 5XFAD and WT cohorts (Figure S4). Analysis of HSV-1 genome copy number in brains of mice at 120 and 144 hours post-inoculation revealed a large variation between animals within a group, as well as a lack of statistically significant difference between WT and 5XFAD mice in copy number (Figure 5B). To summarize, this experiment shows that with respect to the survival rate, pre-existing Aβ plaques do not provide significant protection from acute HSV-1 infection.

**Figure 5.**
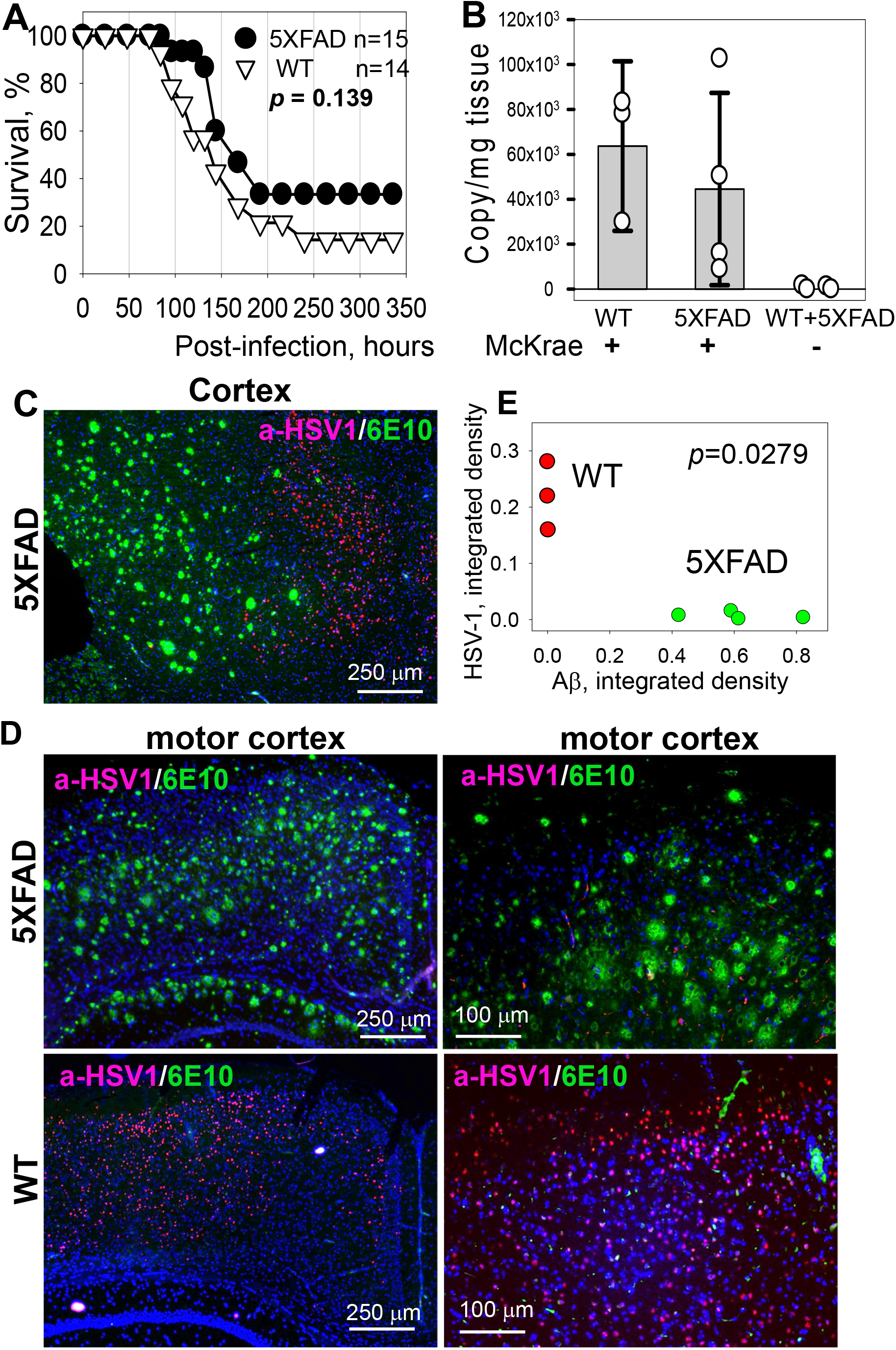
Lack of Protective Effect of Aβ in Aged 5XFAD Mice. (**A**) Survival curves for 7 to 10-month old 5XFAD and WT littermate mice challenged IC with 5×10^3^ PFUs of McKrae per mouse. 5XFAD and WT mice were caged together in random ratios. Statistical significance (*p*) was calculated using log-rank (Mantel-Cox). (**B**) Analysis of the number of HSV-1 genome copies in brains of WT (*n*=3) and 5XFAD (*n*=4) mice IC challenged with McKrae and reached terminal stage of acute encephalitis at 120 and 144 hours post-inoculation. Mean ± SD are shown; n=3 (WT); n=4 (5XFAD). Control group consists of age-matched non-inoculated WT and 5XFAD mice (combined *n*=4). (**C, D**) Co-immunostaining for HSV-1 replication centers (a-HSV1 antibody, red), Aβ plaques (6E10 antibody, green) and nuclei (DAPI, blue) in 7 to 10-month old 5XFAD (*n*=9) and WT littermate (*n*=4) mice, which were challenged IC with 5×10^3^ PFUs of McKrae and succumbed to acute herpes simplex encephalitis within 96-200 hours post-inoculation. (**E**) Analysis of integrated density of HSV-1 replication centers as a function of integrated density of Aβ plaques in the motor cortex of McKrae-inoculated 5XFAD (*n*=4, green) and WT littermates (*n*=3, red). Integrated density of HSV-1 immunostaining signal was found to be different in the motor cortex of WT and 5XFAD mice (\**p_adj_*=0.0279, calculated using ordinary 2-way ANOVA with Šidák’s multiple comparisons test).

### In Aged 5XFAD Mice, HSV-1 Does Not Replicate in Brain Regions with Aβ Plaques

Aged 5XFAD mice showed slightly better survival rate in comparison to the WT littermate cohort (Figure 5A). Difference in the slope of the survival curve seen for these two genotypes suggests differences in response to the HSV-1 infection. Indeed, analysis of HSV-1 replication sites revealed that in 5XFAD mice, HSV-1 infection appears to be suppressed in areas with abundant deposition of Aβ plaques (Figure 5C,D). The most severe effect was observed in motor cortex, the region with a high density of Aβ plaques. While HSV-1 replication sites were abundant in the motor cortex of WT littermates, they were barely detectable in 5XFAD mice (Figure 5D). Quantitative imaging that integrates the signal intensity of HSV-1 replication sites confirmed a statistically significant difference between the densities of HSV-1 replication sites in the motor cortex of 5XFAD and WT littermates (Figure 5E). While Aβ plaques appear to affect distribution of HSV-1 replication centers across brain areas, this phenomenon seems to provide only limited protection to 5XFAD mice, as the difference in survival rates between 5XFADand WT was not significant.

### Lack of Association between HSV-1 and Aβ Plaques in Aged 5XFAD Mice

To test whether the lack of viral replication in areas with a high density of Aβ plaques was due to entrapping of HSV-1 by Aβ plaques, we examined colocalization of Aβ with HSV-1 using coimmunostaining with anti-Aβ antibody H31L21 (green channel) and anti-HSV-1 antibody gB clone T111 (red channel) that was used in previous studies to detect association between HSV-1 and Aβ [7]. Anti-gB detects glycoprotein B of the HSV-1 viral envelope. In aged 5XFAD mice inoculated with McKrae, two out of eight 5XFAD mice challenged with McKrae exhibited a strong signal in the anti-gB channel (Figure 6A, animals #1 and 2), whereas six 5XFAD mice showed weak staining of Aβ plaques by anti-gB (Figure 6A, animal #3). Aged non-infected 5XFAD serving as negative controls showed weak staining of Aβ plaques by anti-gB, which was similar to the intensity observed in six infected 5XFAD animals even after optimization of the staining protocol (Figure 6B). This result suggests that the clone T111 clone cross-reacts with Aβ plaques.

**Figure 6.**
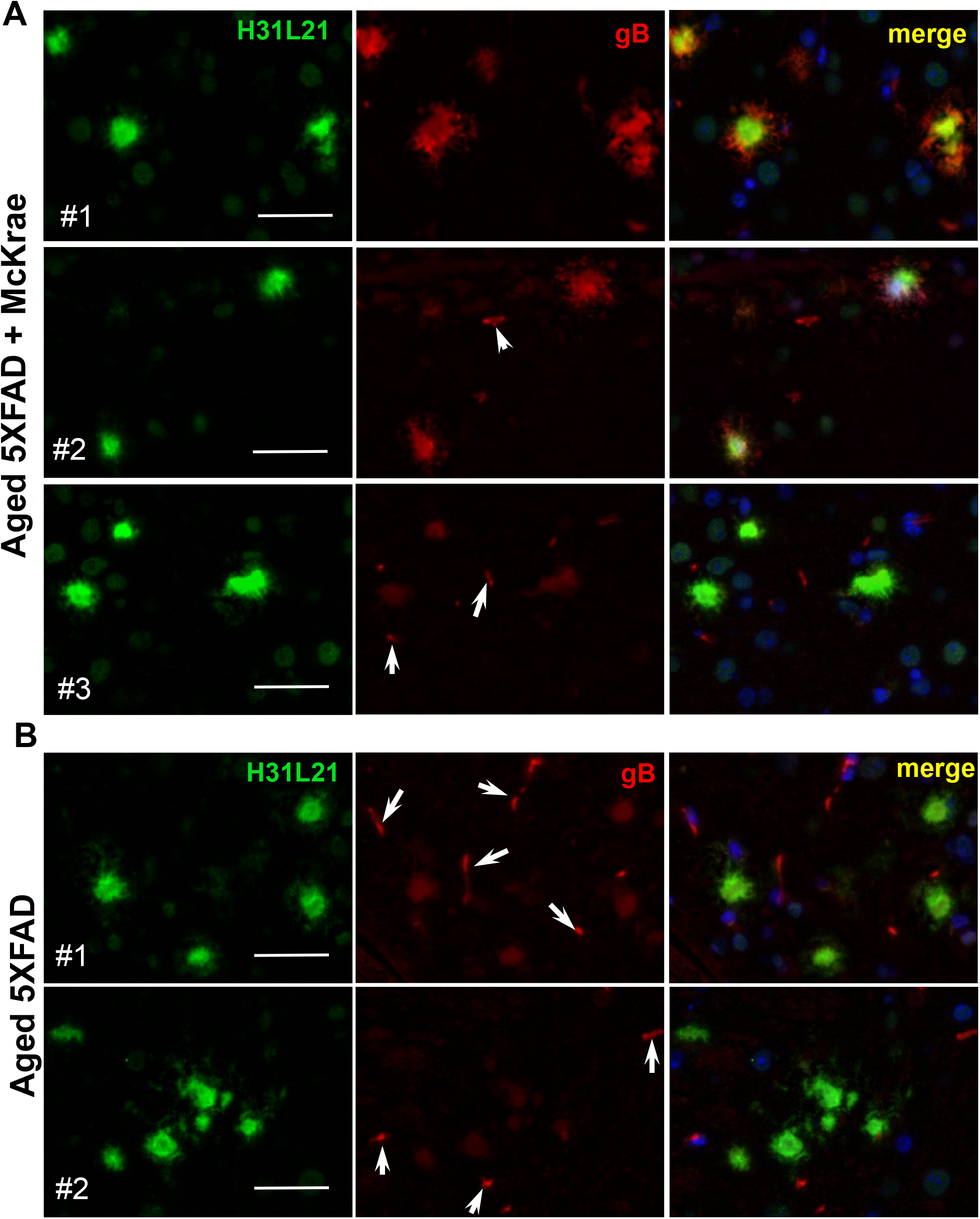
Co-immunostaining for HSV-1 and A β Plaques using anti-gB in Aged 5XFAD Mice. Co-immunostaining for Aβ plaques (H31L21 antibody, green), HSV-1 virus (anti-gB antibody, red) and nuclei (DAPI, blue) in 7 to 10-month old 5XFAD mice, which were challenged IC with 5×10^3^ PFUs of McKrae and succumbed to acute herpes simplex encephalitis within 96-200 hours post-inoculation (*n*=8) (**A**), and uninoculated age-matched 5XFAD (*n*=5) (**B**). Animals #1 and #2 in McKrae-challenged group (**A**) showed strong fluorescence signal intensity and elaborate plaque morphology in the anti-gB channel, whereas animals #3 had the same signal intensity in the anti-gB channel as control group. Arrows point at staining of blood vessels. Scale bars = 20 μm.

To answer the question as to whether the signal in infected 5XFAD mice can be attributed entirely to the cross-reactivity of clone T111 or is, at least in part, due to trapping of HSV-1 by Aβ plaques, we analyzed the degree of colocalization between anti-Aβ and anti-gB antibodies. We reasoned that, if HSV-1 is absent, the cross-reactivity of anti-gB to Aβ plaques should produce a similar degree of colocalization between the two channels in both infected and noninfected brains. If the virus is present in Aβ plaques, the colocalization coefficient between the anti-Aβ and anti-gB channels is expected to be lower in infected versus non-infected mice. For quantitative analysis of colocalization, Manders’ Overlap Coefficients tM1 (for anti-Aβ channel) and tM2 (for anti-gB channels) were calculated for individual plaques from control and experimental 5XFAD groups (Figure S5A,B) [41]. Cross-comparison showed no statistically significant differences between control and experimental groups in tM1 or tM2 values (Figure S5C). The colocalization analysis supports the conclusion that the signal in the infected 5XFAD group was due to cross-reactivity of clone T111 to Aβ plaques.

To further examine whether HSV-1 binds to Aβ plaques, next we used anti-gD and anti-gH antibodies which detect glycoproteins D and H of the HSV-1 envelope, respectively, and do not show cross-reactivity to Aβ plaques. First, co-immunostaining of 5XFAD branes infected with McKrae using a-HSV1 antibody and anti-gD or anti-gH antibodies confirmed that both anti-gD and anti-gH stained the same brain areas and the same cells that were also positive for HSV-1 replication centers (Figure S6). As expected, a-HSV1 detected replication centers and stained nuclei of infected cells, whereas anti-gD or anti-gH labeled the cytoplasm of a-HSV1-positive cells and pericellular spaces in the vicinity of the a-HSV1-positive cells (Figure S6). This experiment confirmed that both anti-gD or anti-gH, can effectively detect the virus. Next, we determined that immunostaining with anti-gD or anti-gH of aged 5XFAD mice inoculated with McKrae reveals the presence of HSV-1 in mouse brain (Figures 7A and S7A). As found previously, multiple brain areas including cortex, hippocampus, thalamus and hypothalamus were affected. However, no signs of colocalization between HSV-1 and Aβ plaques could be found using anti-gD or anti-gH in any of the brain areas (Figures 7A,B and S7A,B). Careful examination of individual Aβ plaques under high magnification showed a total lack of HSV-1 signal in Aβ plaques (Figures 7B and S7B). Imaging of non-infected aged 5XFAD mice confirmed the lack of cross-reactivity of anti-gD or anti-gH to Aβ plaques (Figures 7B and S7B). These data do not support the co-localization of HSV-1 and Aβ plaques.

**Figure 7.**
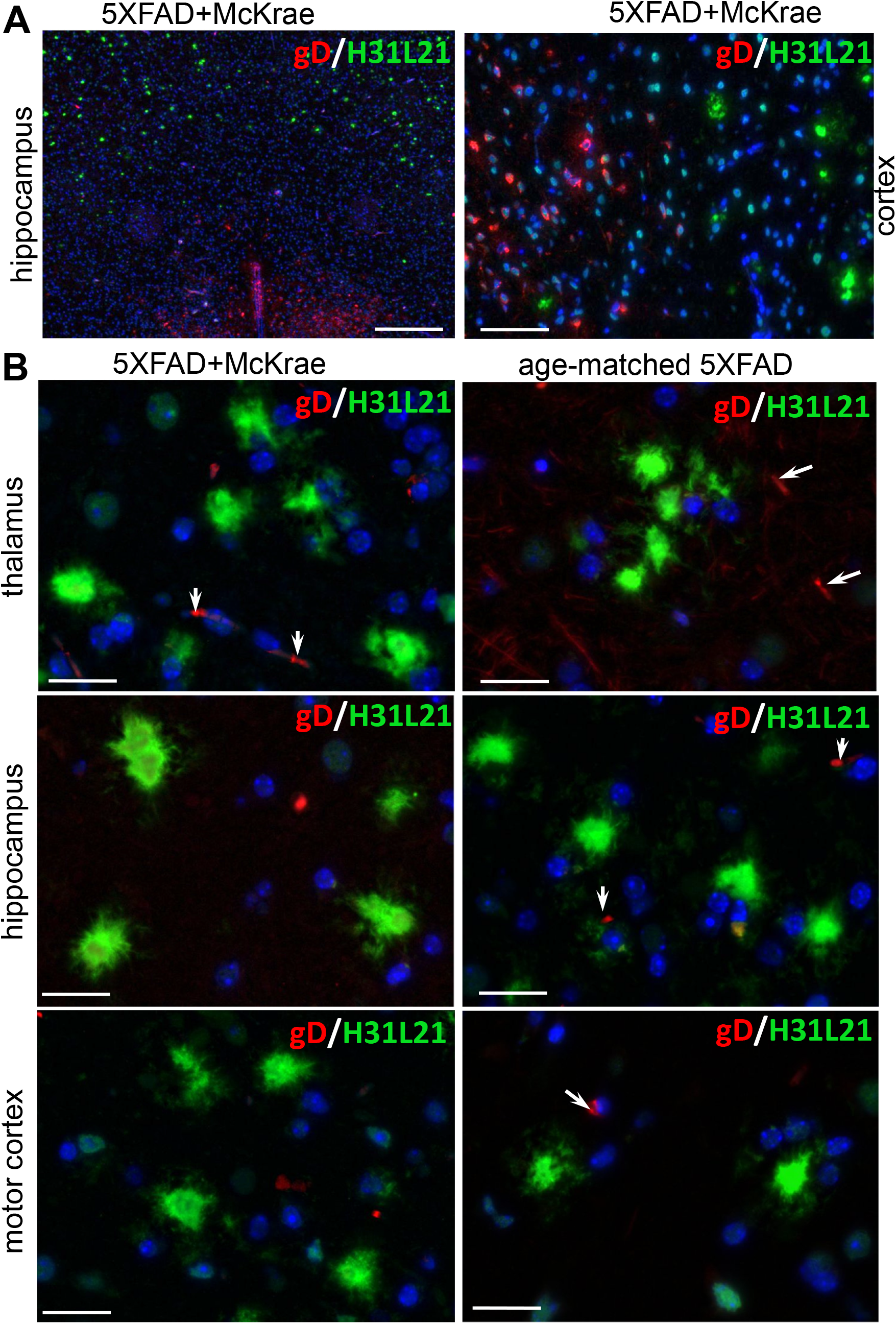
Co-immunostaining for Aβ Plaques and HSV-1 using anti-gD in Aged 5XFAD Mice. Co-immunostaining for Aβ plaques (H31L21 antibody, green), HSV-1 virus (anti-gD antibody, red) and nuclei (DAPI, blue) in infected 7 to 10-month old 5XFAD mice (*n*=8) examined under low (**A**) or high magnifications (**B**), and uninoculated age-matched 5XFAD (*n*=3) (**B**). 5XFAD mice were challenged IC with 5×10^3^ PFUs of McKrae and succumbed to acute herpes simplex encephalitis within 96-200 hours post-inoculation. Arrows point at staining of blood vessels. Scale bars = 100 μm in **A** and 20 μm in **B**.

To explain the absence of HSV-1 in Aβ plaques, we considered the possibility that the virus could have been entrapped by Aβ plaques temporarily, but then quickly cleared by microglia. To test whether HSV-1 was present in Aβ plaques only at the early time points of infection, aged 5XFAD were inoculated with the highest titer of McKrae used in the current work and examined at 24 or 48 hours post-inoculation. Co-immunostaining using anti-Aβ antibody H31L21 and anti-gD antibody did not show any signs of colocalization of HSV-1 with Aβ plaques (Figure S8). We conclude that Aβ plaques do not entrap HSV-1 virus.

## Discussion

Contrary to previous studies [7], the 5XFAD genotype was not protective against acute HSV-1 infection in the current work. This conclusion is supported by experiments that employed two HSV-1 strains, 17syn+ and McKrae, each administered at three different doses into young 5XFAD mice, as well as by the examination of the survival rate of aged 5XFAD mice upon challenge with McKrae. In support of this conclusion, expression of the APP variant associated with familial AD did not alter the region-specific or cell-specific tropisms of HSV-1 strains in 5XFAD mice. These results suggest that the major events in the host-pathogen interaction, such as HSV-1 trafficking, cell invasion or replication, are not affected by APP overexpression. In fact, active sites of HSV-1 replication were found in both APP-positive and negative neurons. Furthermore, contrary to the previous studies [7], young 5XFAD mice infected with HSV-1 did not form Aβ plaques.

The discrepancy between the current and previous studies could be due to the substantially higher dose of HSV-1 used in the previous work [7] in comparison to the current study, or to differences in the strain of HSV-1 used. In the current work, mice were challenged with three doses of McKrae or 17syn+, which were 5-to 10-fold above or below the strain-specific LD_50_ values. An LD_50_ value depends on both the HSV-1 strain and the mouse strain. Previous studies administered 2×10^7^ PFUs per mouse [7], however, it is difficult to estimate how high above the LD_50_ value this dose was, because information regarding the HSV-1 strain was not provided. Thus, there is a possibility that the 5XFAD genotype is indeed protective against HSV-1, yet under a narrower set of experimental conditions, i.e. a specific dose or strain of HSV-1. Another experimental parameters which differed from the previous study was the sex of animals. In the current work, both males and females were used in a random ratio in each experiment, whereas the previous study tested only females [7]. However, analysis of the survival curves for females in the current work also showed a lack of statistically significant differences between 5XFAD and WT cohorts.

Aged 5XFAD animals, in which Aβ plaques are abundant, showed a longer survival and a slightly better survival rate relative to the WT mice, however, the differences between the two groups were not statistically significant (Figure 5A). In aged 5XFAD mice, HSV-1 was found to be excluded from brain areas with abundant Aβ plaques (Figure 5C,D). It is possible that the exclusion effect was responsible for a delay in disease progression in 5XFAD mice suggesting that pre-existing Aβ plaques might provide temporal relief against HSV-1 infection. Nevertheless, because this protective effect occurs only in aged animals, it is not clear that this phenomenon is beneficial evolutionarily, or even whether it reflects an actual biological function of Aβ. From our data, it appears that invasion and replication of HSV-1 in brain areas that lacked Aβ plaques is sufficient for triggering acute simplex herpes virus encephalitis (HSE). Indeed, analysis of the number of viral genome copies revealed no statistically significant differences between old 5XFAD mice and WT littermate controls (Figure 5B).

To test whether the temporally protective effect was due to entrapment of viruses by plaques, colocalization between HSV-1 and Aβ plaques was examined in aged 5XFAD mice that succumbed to acute herpes simplex encephalitis using antibodies to three glycoproteins (gB, gD and gH) of the HSV-1 envelope. Two antibodies (gD and gH) unambiguously showed a lack of colocalization between HSV-1 and Aβ plaques, whereas the results obtained with an anti-gB antibody (clone T111) were not consistent between individual animals within the infected 5XFAD cohort. Co-immunostaining of Aβ plaques in aged 5XFAD mice lacking HSV-1 revealed cross-reactivity of the clone T111 with Aβ plaques. We do not know whether the cross-reactivity of the clone T111 depends on the biological age of a plaque, a plaque-specific Aβ conformation or Aβ_42/40_ ratios within individual plaques or an animal. However, our data indicate that the lack of consistent results with the clone T111 is clearly attributable to its cross-reactivity with Aβ. This cross-reactivity is perhaps not surprising, considering that the gB protein has a segment homologous to the Aβ peptide. To test whether fast clearance by microglia could accounts for the absence of HSV-1 in Aβ plaques, aged 5XFAD animals were examined 24 or 48 hours post- inoculation with HSV-1 using anti-gD antibody. Again, no signs of HSV-1 could be found in Aβ plaques. In summary, with respect to binding of HSV-1 by Aβ plaques, our work arrives at the opposite conclusion from that of Eimer *et al* [7]. Our examination of negative controls, i.e. uninoculated aged 5XFAD mice, was important for arriving at this conclusion.

While the current findings question the anti-viral role of Aβ, this work does not refute the viral hypothesis behind the etiology of late-onset AD. In fact, our observed lack of a protective effect of the 5XFAD genotype (or of Aβ) against acute HSV-1 infection does not report on the potential role of Aβ in latent or chronic HSV-1 infections, which are likely more relevant to the etiology of late-onset AD. Viral or microbial pathogens of the CNS could increase the risk of late-onset AD via multiple, Aβ-dependent or independent mechanisms. Persistent activation of microglia as a result of recurrent reactivation of latent infection or repetitive exposure to viruses might lead to chronic neuroinflammation, which is an important driver of chronic neurodegeneration. It would be interesting to test whether the reactive phenotypes acquired by glia as a result of multiple exposures to HSV-1, or by recurrent reactivation of latent infections, are similar to those observed in late-onset AD disease. Our study did not aim to characterize phenotypic changes in microglia or astrocytes triggered by HSV-1. In fact, we found that astrocytes and microglia were not among the primary cellular targets of HSV-1, at least during the acute stages of infection. However, we did find that microglial cells of ameboid morphology, typical shapes for reactive phonotypes, were seen in sites of active HSV-1 replication, suggesting that microglia respond promptly to HSV-1 invasion. Moreover, the engulfment of HSV-1-infected neurons by reactive microglia we observed suggests that microglia play an active role in fighting the virus by attacking infected neurons. It is plausible to imagine that recurrent reactivation of latent HSV-1 infections over the lifetime would eventually result in substantial neuronal loss to the action of reactive, neurotoxic microglia via a drop-by-drop mechanism.

A series of studies using cultured cells and primary neurons conducted by the Palamara and Grassi laboratories has documented that HSV-1 infection triggers an Aβ-dependent pathological pathway that does not rely on a protective role for Aβ [42–44]. This pathway involves an HSV-1-induced chain of events, which leads to cleavage of APP, accumulation of Aβ peptides along with other APP fragments in nuclei, followed by transcriptional activation of the neprilysin and glycogen synthase kinase 3β (*gsk3β*) genes and impairment of synaptic function [42–44]. Glycogen synthase kinase 3 is one of several kinases that phosphorylates tau [45], a process that is considered to be one of the main driving forces behind AD pathogenesis. This pathway may constitute an alternative mechanism to link viral infection and AD pathology.

A large amount of circumstantial evidence has been put forward in support of the viral hypothesis of AD. Individuals surviving HSE, the acute form of HSV-1 infection of the CNS, show clinical symptoms reminiscent of AD. The brain regions affected in HSE are the same regions compromised in AD [46, 47]. Moreover, in a longitudinal study, individuals positive for anti-HSV-1 IgG and IgM exhibit a higher risk of developing AD relative to control groups [48-50]. Studies using *APOE* knockout (KO), or humanized *APOE3* or *APOE4* transgenic mice reveal that neuroinvasion and spread of HSV-1 in brain is significantly facilitated in *APOE4* mice in comparison to *APOE* KO or *APOE3* mice [51–54]. In support of the viral hypothesis, DNA from HSV-1 colocalizes with Aβ plaques in the brains of AD individuals [55], while HSV-1 is able to seed Aβ fibrils *in vitro* [7]. However, critics of the viral hypothesis argue that viral infections and AD hallmarks often coincide simply because both are common in aged brains, and that infections might be secondary to the development of AD due to a disrupted blood brain barrier. Other studies in support of the viral hypothesis demonstrate that infection of cultured cells or primary neurons with HSV-1 alters APP processing and induces accumulation of Aβ peptides [42, 43, 56–58]. Infecting human cells or mice with HSV-1 can increase the deposition of Aβ [7, 11, 21, 59–61] and of phosphorylated tau [21, 61–64]. However, the lack of a robust cell culture system that can model *latent* HSV-1 infection represents a major obstacle for moving forward to elucidate the role of HSV-1 in AD etiology using cell culture models [65].

Up to 90% of the world population harbors latent HSV-1 infection [66]. How does the viral hypothesis explain that only a relatively small fraction of HSV-1-positive individuals develop late-onset AD? In the vast majority of the human population, HSV-1 infection is latent. The viral hypothesis proposes that the combination of two factors: (i) persistent exposure to pathogens, including recurrent incidents of activation, and (ii) a hyperreactive CNS innate immune system in individuals harboring high-risk variants of *APOE4* and others, is responsible for the etiology of late-onset AD. In agreement with this mechanism, reactivation of HSV-1 during asymptomatic latency in mice was found to upregulate neuroinflammatory response along with phosphorylation and cleavage of tau [62]. Recent animal studies by Palamara and Grassi laboratories provided the most compelling evidence to date linking repeated reactivation of latent HSV-1 infection with AD pathology [21, 67, 68]. Repetitive thermal reactivation of latent HSV-1 infection of the CNS in wild type mice was found to trigger a broad spectrum of neuropathological and clinical changes associated with AD including accumulation of Aβ peptides and Aβ plaques, an increase in phosphorylation of tau, neuroinflammation, oxidative stress, impaired neurogenesis, reactive astrogliosis and memory loss [21, 67, 68]. Establishing HSV-1 latency in neurons is a complex process that is controlled by HSV-1 strain identity on the one hand [69, 70], and a variety of host factors including neuronal subtype, on the other hand [71, 72]. Future studies are needed to establish whether HSV-1 strain identity defines a risk factor for late-onset AD.

Recent epidemiological data offered strong support of a causative link between HSV-1 and late-onset AD. Retrospective studies are based on electronic health records in Taiwan, which enrolled a total of 33,448 subjects, including 8,362 individuals who were newly diagnosed with HSV-1 infection, and 25,086 non-infected controls [73]. The study revealed not only that HSV-1 infection increases the risk of dementia by 2.56-fold at older ages, but also that HSV-1-positive people treated with antiviral drugs were 10-fold less likely to develop dementia later in life, relative to untreated HSV-1-positive individuals [73]. This study establishes that antiviral drugs are very effective as preventative treatments against late-onset AD. Surprisingly, the Einstein Aging Study [74] that monitored rates of dementia incidents between 1993 and 2015 among 70-year or older individuals revealed a gradual decline in dementia incidence in successive birth cohorts. In particular, cohorts born after 1929 have significantly lower rates of dementia development relative to the cohorts born before 1929 upon reaching the same biological age. The same conclusion was reached in the Framingham Heart Study that monitored dementia incidence in individuals from 61 to 101 years old from 1975 until the present time [75]. It is tenable that increased access to antibiotics and antiviral drugs in the second half of the twentieth century is responsible for the decline in the rates of dementia, however, this hypothesis remains to be causally tested.

Using a comprehensive set of experimental conditions which involved inoculation of two HSV-1 strains, each administered at three doses into 5XFAD mice of both sexes, the current work argues against the hypothesis that Aβ exhibits a general anti-viral effect. Aged 5XFAD animals, in which Aβ plaques are abundant, did not exhibit a statistically significant increased survival rate relative to the control mice. Moreover, in aged 5XFAD mice, Aβ plaques were free of HSV-1, suggesting that pre-formed plaques do not entrap the virus. While the current work does not confirm the antiviral role of APP or Aβ, this study does not support nor refutes the hypothesis on the viral etiology of late-onset AD. Alternative hypotheses that do not rely on direct antiviral effects of Aβ should be considered in order to provide experimental support for advancing the viral etiology hypothesis of AD and explain the clear epidemiological findings.

## List of abbreviations

AD: Alzheimer’s Disease
APP: amyloid precursor protein
HHV6: human Herpesviruses 6
HHV7: human Herpesviruses 7
HSE: herpes virus encephalitis
HSV-1: herpes simplex virus 1
IC: intracranial
McKrae: strain of HSV-1
PFUs: plaque forming units
WT: wild type
17syn+: strain of HSV-1

## Declarations

### Ethics approval and consent to participate

The study was carried out in accordance with the recommendations in the Guide for the Care and Use of Laboratory Animals of the National Institutes of Health. The animal protocol was approved by the Institutional Animal Care and Use Committee of the University of Maryland, Baltimore (Assurance Number: A32000-01; Permit Number: 0419007).

### Consent for publication

Not applicable

### Availability of data and materials

All data generated or analyzed during this study are included in this published article and its supplementary information file.

### Competing interests

The authors declare that they have no competing interest.

### Funding

Financial support for this study was provided by National Institute of Health Grants R01 NS045585 and R01 AI128925 to IVB.

### Authors’ contributions

Conceptualization, I.V.B. and O.B.; Methodology, O.B.; Investigation, O.B., K.M. and N.P.P.; Resources, K.M.; Writing – Original Draft, I.V.B.; Writing – Review and Editing, O.B., K.M., and N.P.P.; Supervision, I.V.B. and O.B.; Funding Acquisition, I.V.B.

## Acknowledgments

We thank Natallia Makarava and Iris Lindberg for reading and discussing the manuscript. This work was funded by the NIH (Supplemental awards for R01 NS045585 and R01 AI128925).

## Notes

### Competing Interest Statement

The authors have declared no competing interest.

