## Supplementary material for "Alzheimer’s disease-associated β-Amyloid does not protect against Herpes Simplex Virus 1 brain infection": Figure S1

**Supplemental Figure Legends**

**Figure S1. Dose-Response of Young Female 5XFAD Mouse Model to HSV-1 challenge.** Survival curves for 5-6 weeks old female 5XFAD and wild-type littermate (WT) mice challenged via IC injections with $10^5$, $10^4$ or $10^3$ PFUs of 17syn+ strain per mouse (**A**); or $10^4$, $5x10^3$, or $10^3$ PFUs of McKrae strain per mouse (**B**). 5XFAD and WT littermate mice were caged together in random ratios. Individual plots show independent experiments with number (*n*) of animals of each genotype indicated. Statistical significance (*p*) was calculated using log-rank (Mantel-Cox).

**Figure S2. Region-Specific Tropism of HSV-1 Is Not Altered in 5XFAD Mice. (A)** Immunostaining for HSV-1 replication centers (a-HSV1 antibody, red) and cell nuclei (DAPI, blue) in 5XFAD (*n*=5) and WT littermates (*n*=5) that did not survive IC challenge with $5x10^3$ PFUs of McKrae. **(B)** Immunostaining for HSV-1 replication centers (a-HSV1 antibody, red) and cell nuclei (DAPI, blue) in 5XFAD mice that did not survive IC challenge with $10^4$ PFUs of McKrae, or age-matched 5XFAD mice.

**Figure S3. HSV-1 Does not Induce Formation of Aβ Plaques in Young 5XFAD Mice. (A, B)** Co-immunostaining for Aβ plaques (6E10 antibody, green), HSV-1 replication centers (a-HSV1 antibody, red), and nuclei (DAPI, blue) in 5XFAD mice that survived IC challenge with $10^4$ PFUs of 17syn+ (*n*=3) (**A**) or $10^4$ PFUs of McKrae (*n*=2) (**B**). Animals were examined 2 weeks or 7.5 weeks upon IC challenge. No HSV-1 replication centers could be detected in animals that survived HSV-1 challenge. **(C)** Co-immunostaining for Aβ plaques (6E10 antibody, green), HSV-1 replication centers (a-HSV1 antibody, red), and nuclei (DAPI, blue) in 4.5-month old (left panel, *n*=2) and 11-month old (right panel *n*=2) control 5XFAD mice. In control 5XFAD mice (**C**), 6E10 antibody stains both APP and Aβ plaques (white arrows). However, in HSV-1-inoculated 5XFAD mice (**A, B**), only intracellular APP staining can be seen.

**Figure S4. Lack of Protective Effect of Aβ in Aged Female 5XFAD Mice.** (**A**) Survival curves for 7 to10- month old female 5XFAD and WT littermate mice challenged IC with $5x10^3$ PFUs of McKrae per mouse. 5XFAD and WT mice were caged together in random ratios. Statistical significance (*p*) was calculated using log-rank (Mantel-Cox).

**Figure S5. Analysis Manders' Overlap Coefficients in Aged 5XFAD Mice. (A, B)** Distribution of tM1 (green channel, Aβ) and tM2 (red channel, gB) colocalization coefficients in 5XFAD mice challenged IC with $5x10^3$ PFUs McKrae ($n$=8) (**A**) and age-matched 5XFAD mice ($n$=5) (**B**). Individual animals are color-coded; each circle represents an individual Aβ plaque. (**C**) Significance testing of colocalization data. The box-and-whisker plot of tM1 and tM2 colocalization coefficients in experimental and controls 5XFAD groups are shown in panels **A** and **B**, respectively. The midline of the box-and-whisker plot denotes the median, the x represents the mean, and the ends of the box plot denote the $25^{th}$ and $75^{th}$ percentiles.

**Figure S6. Antibodies to Viral Replication Sites and Envelop Proteins gD and gH Stain the Same Cells and Brain Regions.** Co-immunostaining for HSV-1 replication centers (a-HSV1 antibody, green) and viral envelope proteins gD or gH (anti-gD or anti-gH antibodies, red) in 6-week old 5XFAD mice ($n$=6) infected with $10^4$ PFUs of McKrae.

**Figure S7. Co-immunostaining for Aβ Plaques and HSV-1 using anti-gH in Aged 5XFAD Mice.** Co-immunostaining for Aβ plaques (H31L21 antibody, green), HSV-1 virus (anti-gH antibody, red) and nuclei (DAPI, blue) in infected 7 to 10-month old 5XFAD mice ($n$=8) examined under low (**A**) or high magnifications (**B**), and uninoculated age-matched 5XFAD ($n$=3) (**B**). 5XFAD mice were challenged IC with $5x10^3$ PFUs of McKrae and succumbed to acute herpes simplex encephalitis within 96-200 hours post-inoculation. Arrows point at staining of blood vessels. Scale bars = 100 μm in **A** and 20 μm in **B**.

**Figure S8. Co-immunostaining for Aβ Plaques and HSV-1 using anti-gD in Aged 5XFAD Mice.** Co-immunostaining for Aβ plaques (H31L21 antibody, green), HSV-1 virus (anti-gD antibody, red) and nuclei (DAPI, blue) in 10-month old 5XFAD mice challenged IC with $10^4$ PFUs of McKrae examined at 24 hours ($n$=5) or 48 hours ($n$=5) post-inoculation. Arrows point at staining of blood vessels.

#### Figure S1

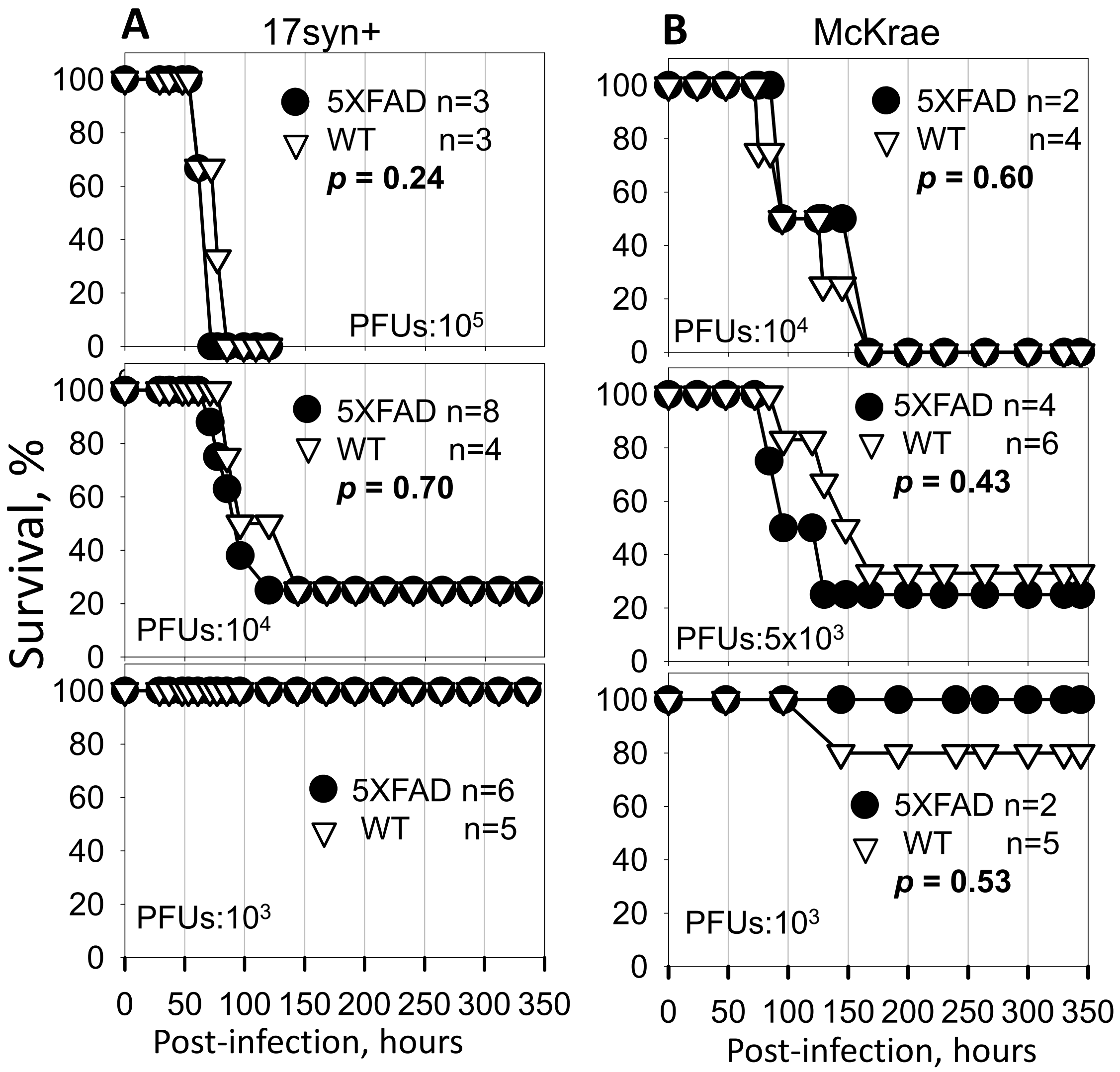
A
17syn+
B
McKrae
Survival, %
5XFAD n=3
WT n=3
p = 0.24
PFUs:10^5
5XFAD n=8
WT n=4
p = 0.70
PFUs:10^4
5XFAD n=6
WT n=5
PFUs:10^3
5XFAD n=2
WT n=4
p = 0.60
PFUs:10^4
5XFAD n=4
WT n=6
p = 0.43
PFUs:5x10^3
5XFAD n=2
WT n=5
p = 0.53
PFUs:10^3
Post-infection, hours
Post-infection, hours

**Figure S2**

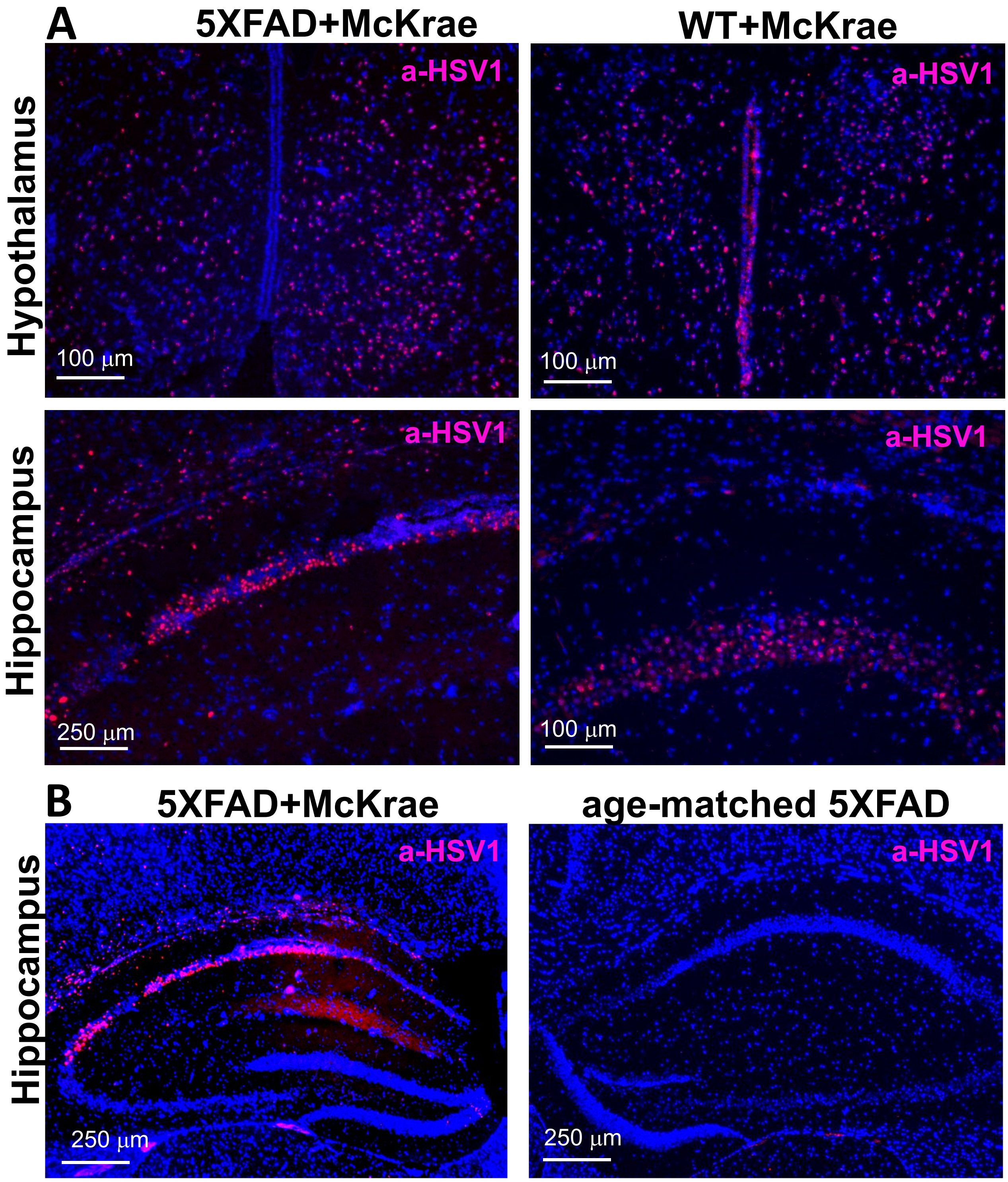

A
5XFAD+McKrae
WT+McKrae
Hypothalamus
a-HSV1
a-HSV1
100 µm
100 µm
Hippocampus
a-HSV1
a-HSV1
250 µm
100 µm
B
5XFAD+McKrae
age-matched 5XFAD
Hippocampus
a-HSV1
a-HSV1
250 µm
250 µm

#### Figure S3

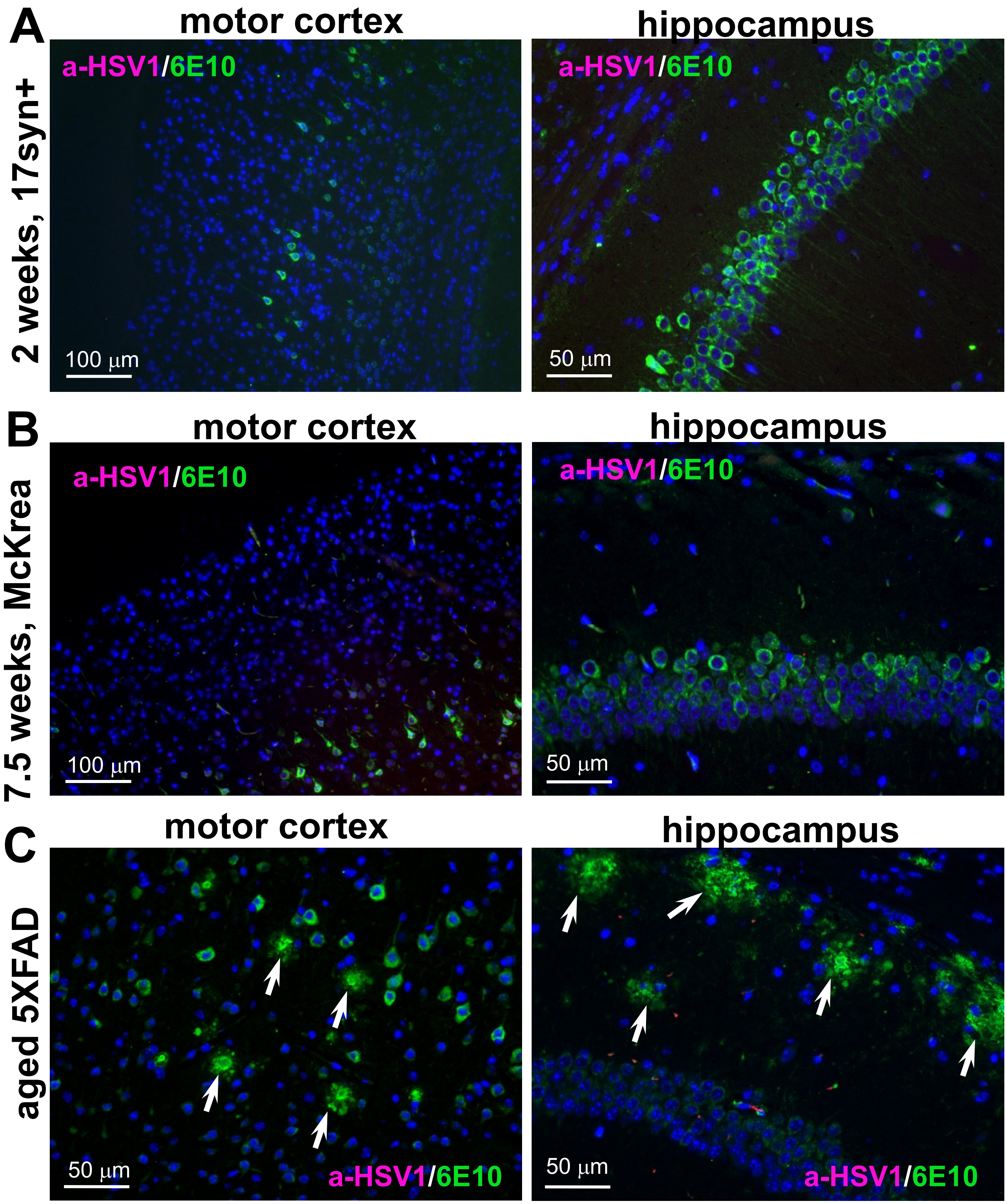
A
motor cortex
hippocampus
2 weeks, 17syn+
a-HSV1/6E10
a-HSV1/6E10
100 µm
50 µm
B
motor cortex
hippocampus
7.5 weeks, McKrea
a-HSV1/6E10
a-HSV1/6E10
100 µm
50 µm
C
motor cortex
hippocampus
aged 5XFAD
50 µm
a-HSV1/6E10
50 µm
a-HSV1/6E10

**Figure S4**

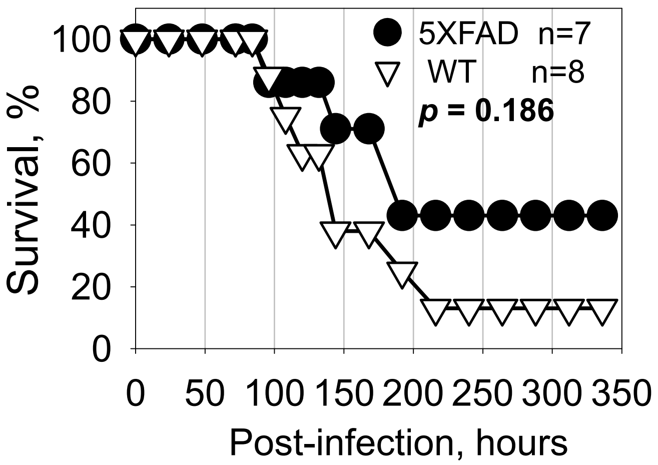

5XFAD n=7
WT n=8
p = 0.186
Survival, %
0
20
40
60
80
100
0
50
100
150
200
250
300
350
Post-infection, hours

### Figure S5

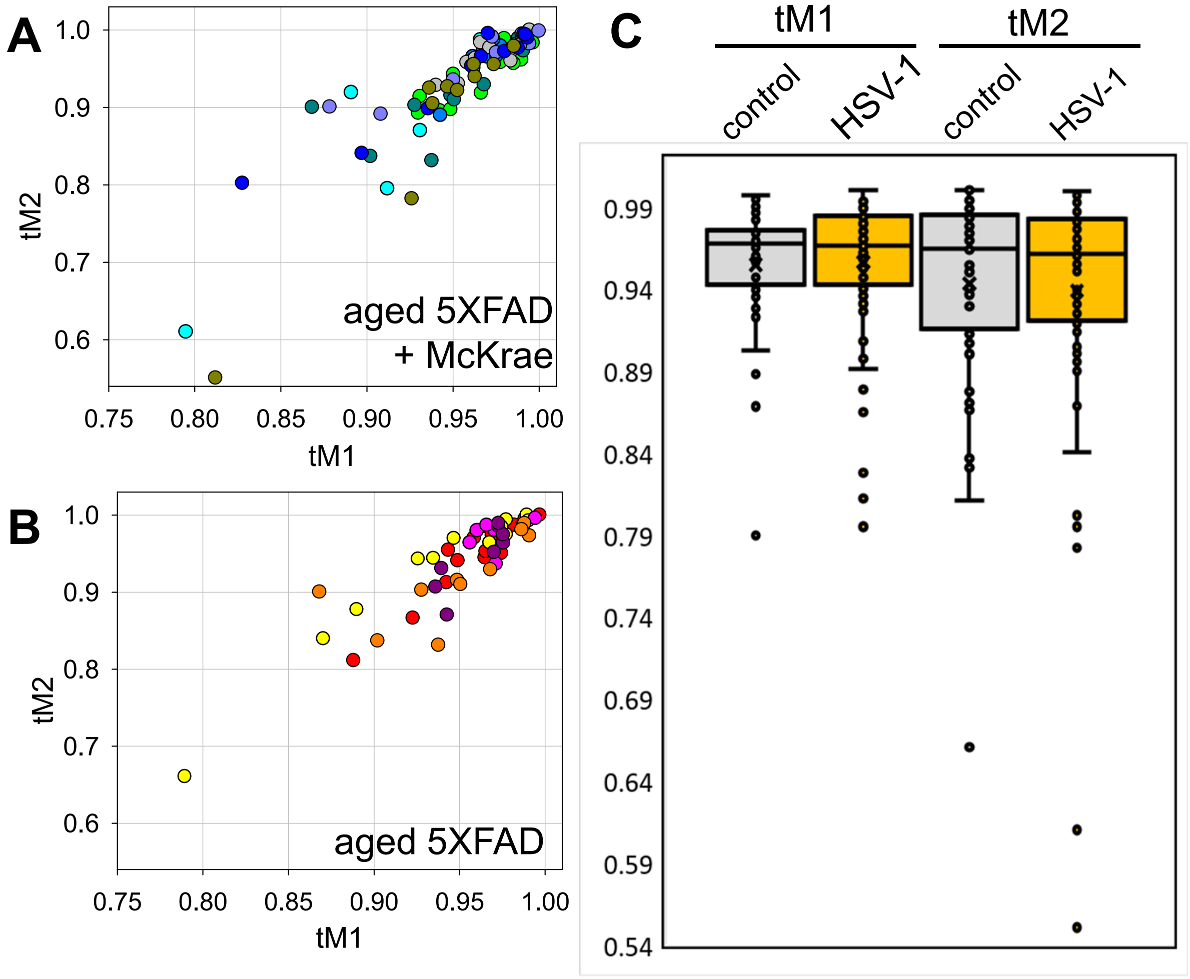
A
1.0
0.9
0.8
0.7
0.6
tM2
aged 5XFAD
+ McKrae
0.75
0.80
0.85
0.90
0.95
1.00
tM1
B
1.0
0.9
0.8
0.7
0.6
tM2
aged 5XFAD
0.75
0.80
0.85
0.90
0.95
1.00
tM1
C
tM1
tM2
control
HSV-1
control
HSV-1
0.99
0.94
0.89
0.84
0.79
0.74
0.69
0.64
0.59
0.54

### Figure S6

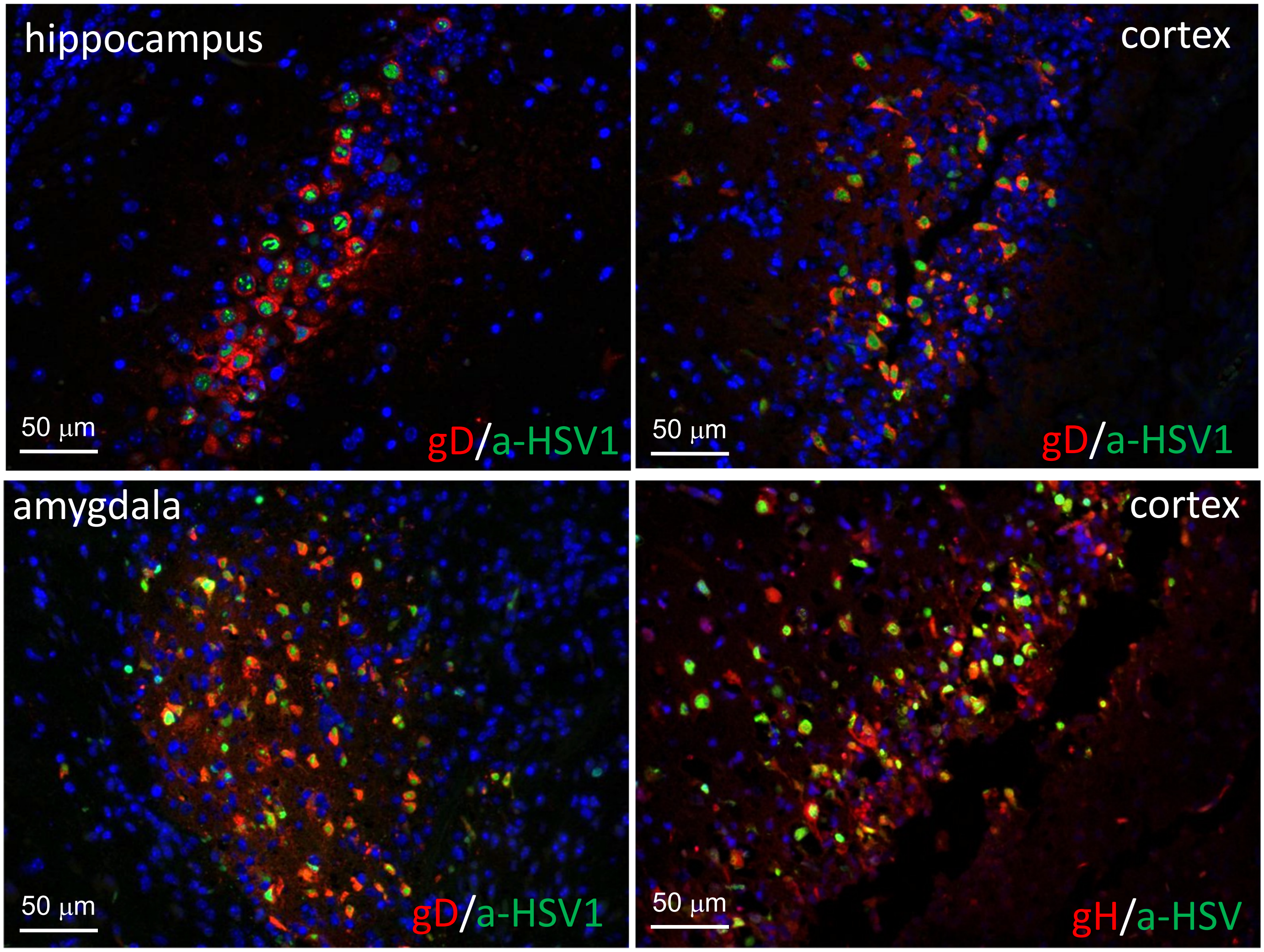
hippocampus
50 μm
gD/a-HSV1
cortex
50 μm
gD/a-HSV1
amygdala
50 μm
gD/a-HSV1
cortex
50 μm
gH/a-HSV

**Figure S7**

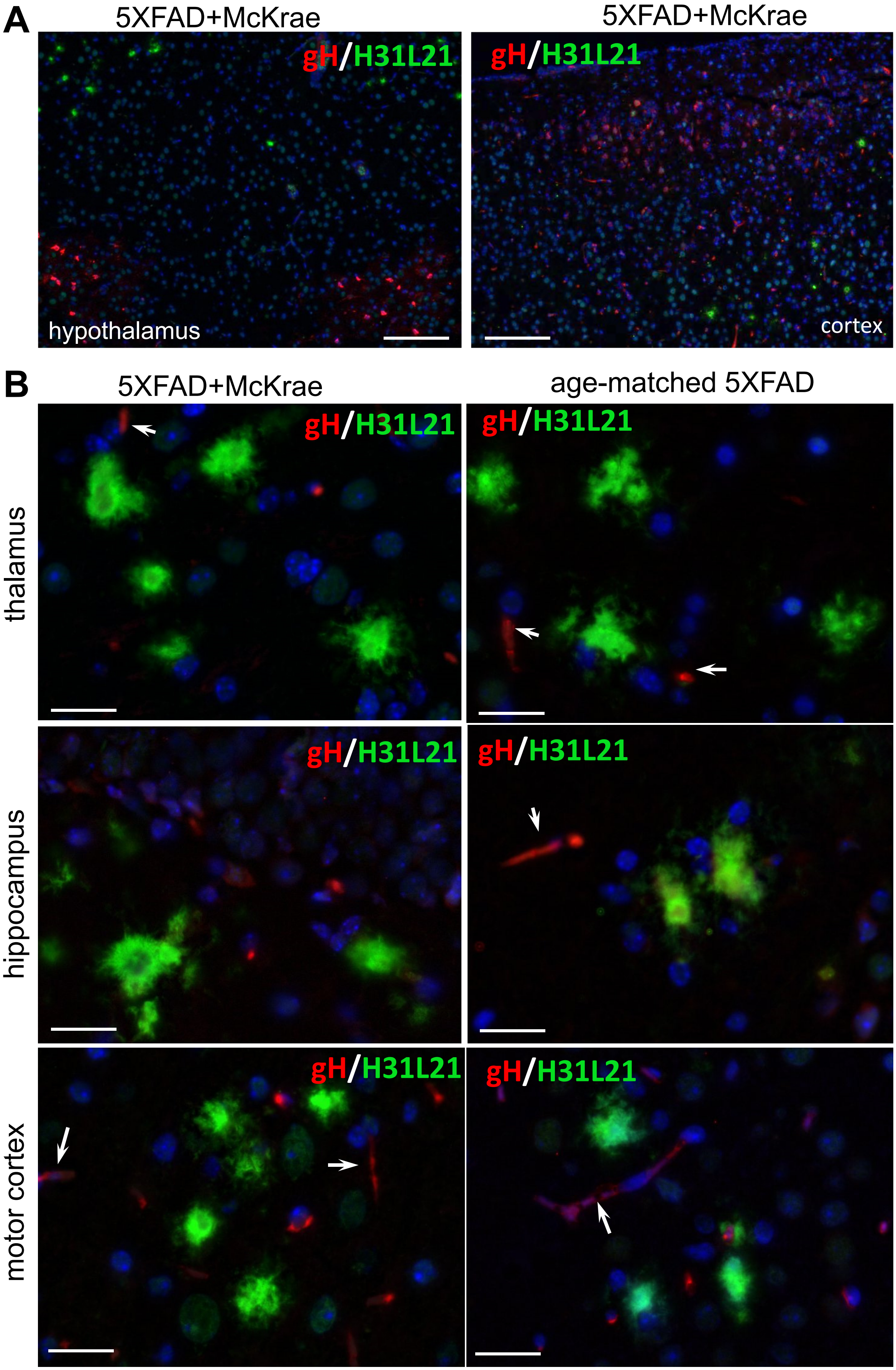
A
5XFAD+McKrae
5XFAD+McKrae
gH/H31L21
gH/H31L21
hypothalamus
cortex
B
5XFAD+McKrae
age-matched 5XFAD
thalamus
hippocampus
motor cortex
gH/H31L21
gH/H31L21
gH/H31L21
gH/H31L21
gH/H31L21
gH/H31L21

### Figure S8

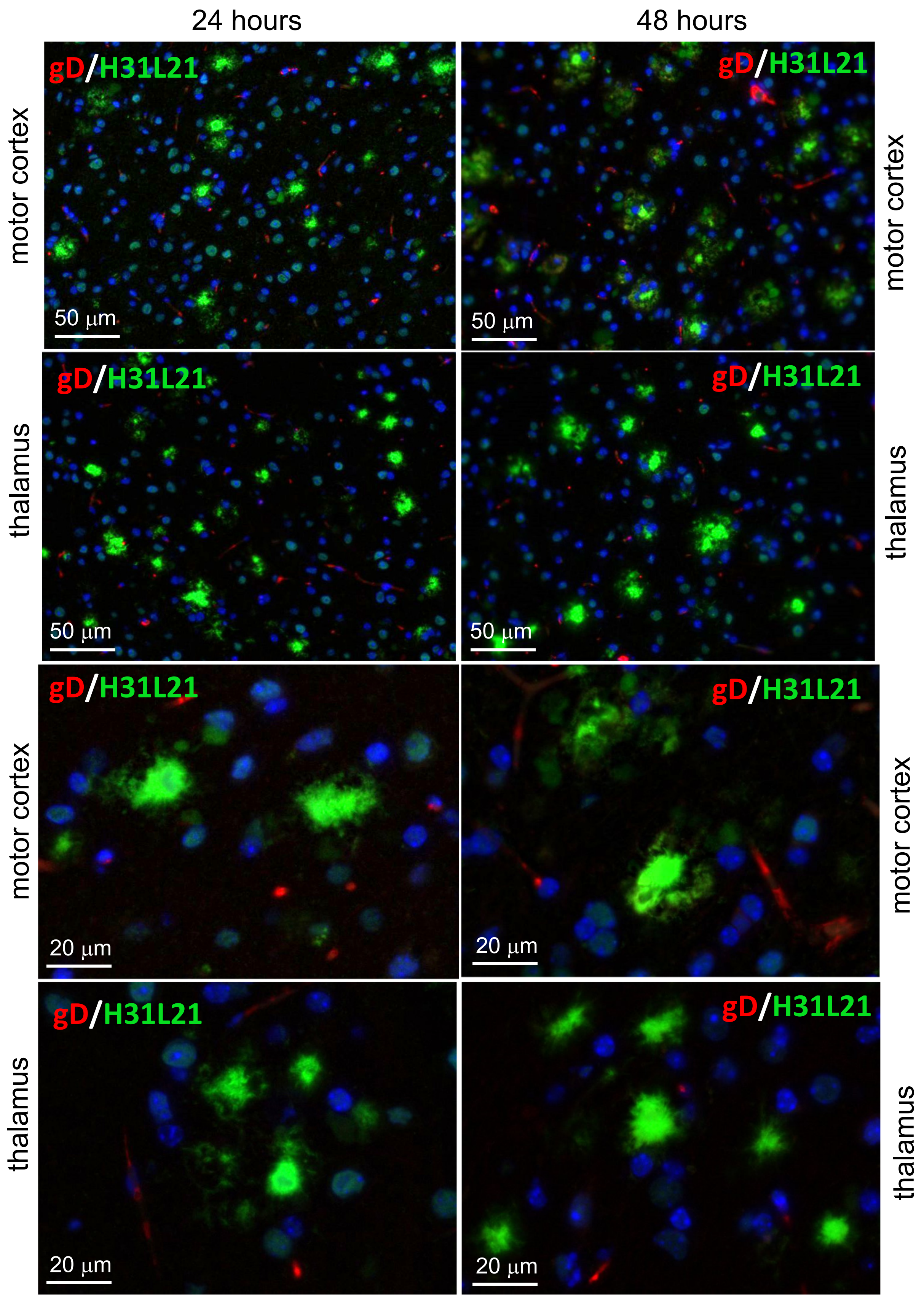
24 hours
48 hours
motor cortex
thalamus
motor cortex
thalamus
gD/H31L21
50 μm
20 μm
